# Vacuolar type H^+^ ATPase is involved in stress responses in *Leishmania mexicana* by regulating the lysosomal pH

**DOI:** 10.64898/2026.08.21.746154

**Authors:** Çağla Alagöz, Andreia Albuquerque-Wendt, Eva Gluenz

**Author notes:** Department of Parasitology, Faculty of Science, Charles University, Prague, Czech Republic Parasite Chemotherapy Unit, Swiss Tropical and Public Health Institute, Allschwil, Switzerland University of Basel, Basel, Switzerland; Global Health and Tropical Medicine, GHTM, LA-REAL, Instituto de Higiene e Medicina Tropical, IHMT, Universidade NOVA de Lisboa, Lisboa, Portugal.

## Abstract

Vacuolar H^+^ ATPases (v-ATPases) are conserved proton pumps that support diverse biological functions through acidification of cellular organelles. The protozoan parasite *Leishmania* requires its v-ATPase for survival in the sand fly vector and mammalian host, but genetic mutants remain viable *in vitro*. To gain further insight into this conditionally lethal phenotype, we first mapped organellar localization of the v-ATPase by co-localisation imaging of fluorescently tagged v-ATPase subunits and organelle markers. The v-ATPase signal was strongest in the flagellar pocket region, consistent with enrichment in the contractile vacuole complex (CVC). To define the conditions that require a functional v-ATPase, deletion mutants were exposed to different stresses (pH, temperature, osmolality, dense culture). All tested deviations from standard culture conditions affected the mutants’ growth rate, viability or both. Despite differences in phenotype severity, all stressors triggered the formation of a large autolysosome, positive for the autophagy marker protein ATG8 and the lysosomal enzyme cysteine peptidase A, indicating an arrest at the final step of autophagy. Measurements with the pH sensor pHLuorin2 showed that the luminal pH of the lysosomes was 5.6 in unperturbed promastigotes and 7.1 in v-ATPase mutants. These data support a canonical function for the *Leishmania* v-ATPase in lysosome acidification and autophagy, which is essential for parasite differentiation, and identify the poorly characterized *Leishmania* CVC as another major site of v-ATPase concentration.

## Introduction

Regulation of intracellular and organellar pH is essential for many biological processes in eukaryotic organisms. While the majority of subcellular compartments have a neutral pH under physiological conditions, some organelles such as lysosomes and endosomes possess an acidic pH for optimal enzyme activity and cellular functions. Vacuolar-type H^+^ ATPases (henceforth v-ATPases) are important evolutionarily conserved proton pumps responsible for the acidification of organellar lumens across eukaryotic life forms (Mulkidjanian et al., 2007). v-ATPases are large multi-subunit protein complexes that are composed of a membrane-associated V_o_ domain and a peripheral V_1_ domain (Vasanthakumar and Rubinstein, 2020). The energy derived from ATP hydrolysis by the catalytic V_1_ domain is utilized by the V_o_ domain for rotation and pumping protons across biological membranes (Imamura et al., 2003). The v-ATPase complex contains a total of at least 14 different subunits, some of which are found in multiple copy numbers (Forgac, 2007). In yeast v-ATPase, subunits A, B, C, D, E, F, G and H constitute the V_1_ domain, whereas V_o_ domain is composed of subunits a, d, e, f, c_8_, c’ and c” (Wang et al., 2023; Wang et al., 2020a). Although overall structure and function of the v-ATPases are well conserved across eukaryotes, some differences in subunit composition and/or copy number are known. The subunit c’ of the c-ring, for example, is not present in the mammalian v-ATPases. In the V_o_ domain of the mammalian v-ATPases, on the other hand, subunit Ac45 and prorenin receptor (PRR) are found, which are absent from yeast v-ATPases (Wang et al., 2020a). The most striking diversification was found in the protist *Paramaecium tetraurelia*, with 17 distinct V_o_a subunits (Wassmer et al., 2006).

v-ATPases localise to the membrane of a range of intracellular organelles including lysosomes, endosomes, Golgi-derived vesicles in mammalian systems and to the lysosome-like vacuoles of yeast and plants (Kane, 2006). In protozoan parasites, v-ATPases are also found in specialized organelles such as the plant-like vacuole of *Toxoplasma gondii*, the contractile vacuole complex (CVC) and acidocalcisomes of *Trypanosoma cruzi*, and acidocalcisomes in *Trypanosoma brucei* (Docampo et al., 1995; Huang et al., 2014; Stasic et al., 2019; Ulrich et al., 2011; Vercesi et al., 1994). The function of v-ATPase in regulating organellar pH is intimately connected to essential biological processes, including endocytosis, intracellular trafficking, protein sorting, as well as autophagy (Collins and Forgac, 2020; Forgac, 2007; Kane, 2006; Yoshimori et al., 1991).

Macroautophagy, hereafter autophagy, is a cellular process whereby proteins and larger cellular components are enveloped in a membranous structure called the autophagosome and, after fusion with a lysosome, degraded and recycled to the cytoplasm (Mizushima, 2007). While autophagy is upregulated in response to various stress signals including starvation, hypoxia and damaged organelles, it is a continuous process at basal level, ensuring cellular homeostasis and quality control (Mizushima, 2007; Mizushima, 2009). When autophagy is induced in the cells, an organelle called phagophore or isolation membrane is formed *de novo* (Mizushima, 2007). Eventually the phagophore structure matures into a double-membrane enclosed autophagosome, engulfing the cytosolic material destined for degradation (Mizushima, 2007). Autophagosomes are delivered to lysosomes for the subsequent degradation of the autophagic material by lysosomal enzymes inside this newly formed autolysosome structure (Mizushima, 2007). Numerous autophagy-related (ATG) proteins with crucial roles in these processes were first identified in yeast mutant screens (Tsukada and Ohsumi, 1993) and are conserved across most eukaryotes (Zhang and Mizushima, 2023), including in protozoan parasites of the family Trypanosomatidae (clade Kinetoplastea) (Brennand et al., 2012; Sakamoto et al., 2021; Williams et al., 2006). Parasites of the genus *Leishmania spp*. cause the neglected tropical disease Leishmaniasis. *Leishmania* have a digenetic life cycle, where they alternate between a mammalian host and an insect vector. Female phlebotomine sandflies transmit the promastigote form of the parasites to a mammalian host during a blood meal. The promastigotes deposited on the skin of the mammalian host are taken up by phagocytic cells, mainly macrophages. Inside the macrophages, changes in temperature (from 27°C to 34-37°C) and pH shift (from neutral to acidic) trigger differentiation of promastigotes into amastigote forms. This differentiation involves extensive remodeling of organelles and cellular morphology (Cull et al., 2014; Wheeler et al., 2016). The parasite cell body becomes rounded, and the long external flagellum gets reduced greatly in size (Wheeler et al., 2015). The study of a dominant negative *L. major* VPS4 mutant first established a link between autophagy and differentiation (Besteiro et al., 2006).

Further mechanistic studies in *L. mexicana* showed that mutants lacking the cysteine peptidases CPA and CPB were defective in autophagy and metacyclogenesis (Williams et al., 2006), that ATG5 is essential for autophagosome formation (Williams et al., 2012) and at least one ATG4 cysteine peptidase is required for autophagy (Williams et al., 2013). These studies established that autophagy is a relevant and essential process for completion of the *Leishmania* life cycle and showed that while some key proteins and mechanisms are widely conserved, there are also lineage-specific losses and diversification of protein repertoires and functions (review (Romano et al., 2023).

v-ATPases are generally involved in the final step of autophagy, by acidifying lysosomal pH and hence providing an optimal environment for many lysosomal hydrolases to function (Nakamura et al., 1997; Yoshimori et al., 1991). Treatment of *Leishmania* with the v-ATPase inhibitor Bafilomycin A1 (BafA1) resulted in perturbed autophagy (Williams et al., 2006). In a systematic gene deletion screen of the transporter proteins of *Leishmania mexicana*, we identified v-ATPase as an essential protein complex for parasites’ survival in the mammalian host, as well as for sandfly colonization and metacyclogenesis (Albuquerque-Wendt et al., 2025; Sadlova et al., 2026). To dissect the physiological functions of the *Leishmania* v-ATPase and define its role in autophagy, we studied here the response of v-ATPase deletion mutants to different environmental conditions encountered by the parasites over the course of their life cycle. We found that loss of v-ATPase function, through genetic ablation of one of its subunits, or by pharmacological inhibition results in the alkalinisation of the lysosome and Atg5-dependent formation of large autolysosomes that cannot be resolved, indicating a stalling at the last step of autophagy.

## Results

### *v-ATPase knockout mutant promastigotes are sensitive to acidic and alkaline pH, elevated temperature and hyperosmotic stress* in vitro

v-ATPases are known to be involved in organellar pH homeostasis, by localizing to the membrane of intracellular organelles and pumping protons across the membranes. Our previous work revealed that the v-ATPase complex is dispensable for *L. mexicana* promastigote growth *in vitro* under standard cell culture conditions (pH 7.4, 27°C) but the parasites’ growth is impaired in pH 5.5 cell culture medium (Albuquerque-Wendt et al., 2025). Expressing an episomal copy of the gene encoding for v-ATPase subunit V_1_E (LmxM.36.3000) in a V_1_E knockout background (V_1_E AB) rescued this growth phenotype in acidic pH (Figure 1A).

**Figure 1:**
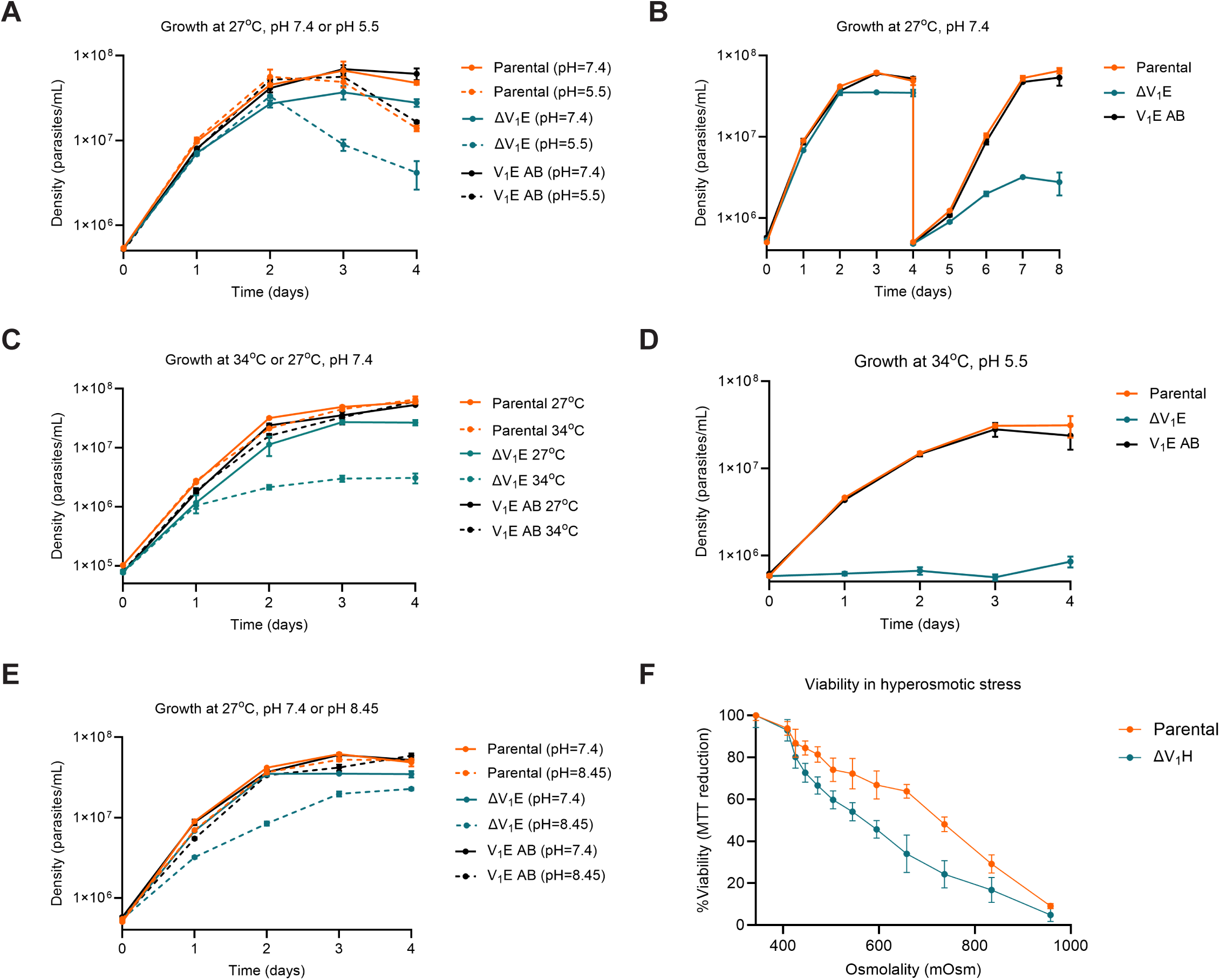
v-ATPase knockout mutant promastigotes are sensitive to stress conditions *in vitro*. (**A**) Continuous growth of v-ATPase ΔV_1_E (teal), V_1_E addback (AB, black) and parental (orange) promastigotes in standard M199 medium at pH 7.4 (solid lines) and in M199 medium pH adjusted to 5.5 (dashed lines). (**B**) Growth of ΔV_1_E, V_1_E AB and parental cell lines in standard M199 medium at pH 7.4. After 4 days of continuous growth, cultures were diluted back to 5x10^5^ cells/mL for an additional 4 days of growth. (**C**) Growth profiles of ΔV_1_E, V_1_E AB and parental cell lines in standard M199 medium, at pH 7.4 and at 27°C (solid lines) or 34°C (dashed lines). (**D**) Continuous growth profiles as in (C), pH of M199 medium adjusted to pH 5.5 for axenic amastigote differentiation. (**E**) Continuous growth profiles of v-ATPase ΔV_1_E, V_1_E AB and parental promastigotes as in (A), pH of M199 medium adjusted to 8.45 (dashed lines). Note, the cultures at pH 7.4 shown here are the same measurements as days 1-4 in(B). Each point in the growth curves represents average cell density measurements from three replicate cultures. Error bars show standard deviation. (**F**) MTT assay measuring the viability of ΔV_1_H (teal) and parental (orange) promastigotes in hyperosmotic stress. Osmolality of the M199 medium was adjusted by addition of sorbitol. Parasites were incubated in hyperosmotic conditions for 24 hours. Each point in the curve represents average viability calculation from 6-8 replicates. Error bars indicate standard deviation.

We noted that during their logarithmic phase of growth, ΔV_1_E parasites grew comparable to parental and V_1_E AB controls. However, as the parasites approach stationary phase, ΔV_1_E promastigotes grew slower than parental and AB cells (Figure 1A). To investigate the growth profiles of parasites after having reached stationary phase, we diluted the promastigotes grown in neutral pH for four continuous days back to their initial density and recorded their growth for an additional four days (Figure 1B). This showed that ΔV_1_E cells recovered poorly from having been in a stationary phase culture, growing more slowly and reaching a plateau early, whereas parental and V_1_E AB parasites recovered their normal rate of growth after one day of lag-phase (Figure 1B). This indicated that the ΔV_1_E promastigotes may be sensitive to changes in nutrient availability in dense cultures.

*L. mexicana* causes cutaneous leishmaniasis, where the parasites reside in the phagolysosomal compartments of the macrophages at the skin. Therefore, the parasites experience a temperature shift from 20-28°C in the sandflies to >30°C in the skin of the vertebrate host. To test whether the ΔV_1_E mutant could tolerate changes in temperature, we tested the growth of ΔV_1_E promastigotes at 34°C in neutral M199 medium and in acidic pH conditions. ΔV_1_E parasites were sensitive to elevated temperature in both pH conditions (Figure 1C, D). In neutral pH, parasites proliferated in the first 24 hours, but their growth stalled after ca. 3-4 cell doublings, reaching a plateau at a cell density 10-fold lower compared to the ΔV_1_E promastigotes grown at 27°C (Figure 1C). At pH 5.5, incubation at 34°C completely prevented growth of the ΔV_1_E parasites (Figure 1D).

We recently reported that knockout of v-ATPase subunits impaired the parasites’ ability to colonise and survive in *Lu. longipalpis* sandflies (Sadlova et al., 2026). Throughout their development in the sandfly, *Leishmania* are exposed to a varied environment, including changes in pH from around 9 in the anterior midgut to 6.5-7.0 in the posterior midgut (Fazito do Vale et al., 2007). To test whether v-ATPase knockout promastigotes can tolerate alkaline pH *in vitro*, the parental control line, ΔV_1_E and V_1_E AB promastigotes were grown in standard M199 medium at pH 7.4 and in M199 medium adjusted to pH 8.45. ΔV_1_E parasites grew slower in alkaline pH compared to parental and V_1_E AB controls (Figure 1E). This indicates that the v-ATPase complex is needed not only for parasites’ adjustment to acidic pH, but also in alkaline conditions.

Another environmental variable encountered between the midgut of the sandflies and the phagolysosome of macrophages is likely osmolality. v-ATPase function is required for osmoregulation in *Naegleria* and *Paramecium*, where treatment with v-ATPase inhibitors BafA1 or concanamycin leads to impairment of water flow in or out of the cell (Allen and Naitoh, 2002; Velle et al., 2023). To test whether the *L. mexicana* v-ATPase was needed for the parasites to withstand hyperosmotic stress, the v-ATPase subunit H knockout (ΔV_1_H, gene ID: LmxM.21.1340) and parental controls were subjected to 340-960 mOsm medium for 24 hours. MTT viability assays indicated that ΔV_1_H promastigotes were more sensitive to hyperosmotic stress (Figure 1F) compared to the parental cells, implying that the v-ATPase is involved in osmoregulation in *Leishmania*. This is consistent with the localization of *L. mexicana* v-ATPase subunits V_1_H and V_1_G in the vicinity of the flagellar pocket in *L. mexicana* (Albuquerque-Wendt et al., 2025) where the CVC is located (Hair et al., 2024).

### Subcellular localization of the v-ATPase

Here, we extended the analysis of subcellular localisations to the other v-ATPase subunits, to investigate whether localization was consistent across all subunits of the protein complex. We successfully tagged 12 out of 14 subunits of the v-ATPase (including isoform variants) at either N- or C-terminus, or both (Supplementary table 1). Two of the subunits constituting the c-ring (c and c’, LmxM.21.1800, LmxM.23.0130) could not be tagged at either of the termini. We observed that the signal in the vicinity of the flagellar pocket was consistent for all tagged subunits, both in promastigotes and axenic amastigotes (Supplementary figure 1A-F). In addition, imaging of a double-tagged cell line with subunit d from the V_o_ domain (LmxM.05.1140) tagged with mNeonGreen (mNG) on its C-terminus and subunit V_1_E tagged with mCherry (mCh) on its N-terminus showed perfect overlap of the two proteins (Supplementary figure 2A).

To increase the resolution of imaging and obtain more information about the subcellular localization of the v-ATPase, we employed ultrastructure expansion microscopy (U-ExM) to image subunit V_1_H tagged with 3x myc epitope at the N-terminus. Individual myc::V_1_H puncta were concentrated near the flagellar pocket; instead of the diffuse cell body signal observed in live cell imaging, we saw here additional puncta in the cytosol (Figure 2A).

**Figure 2:**
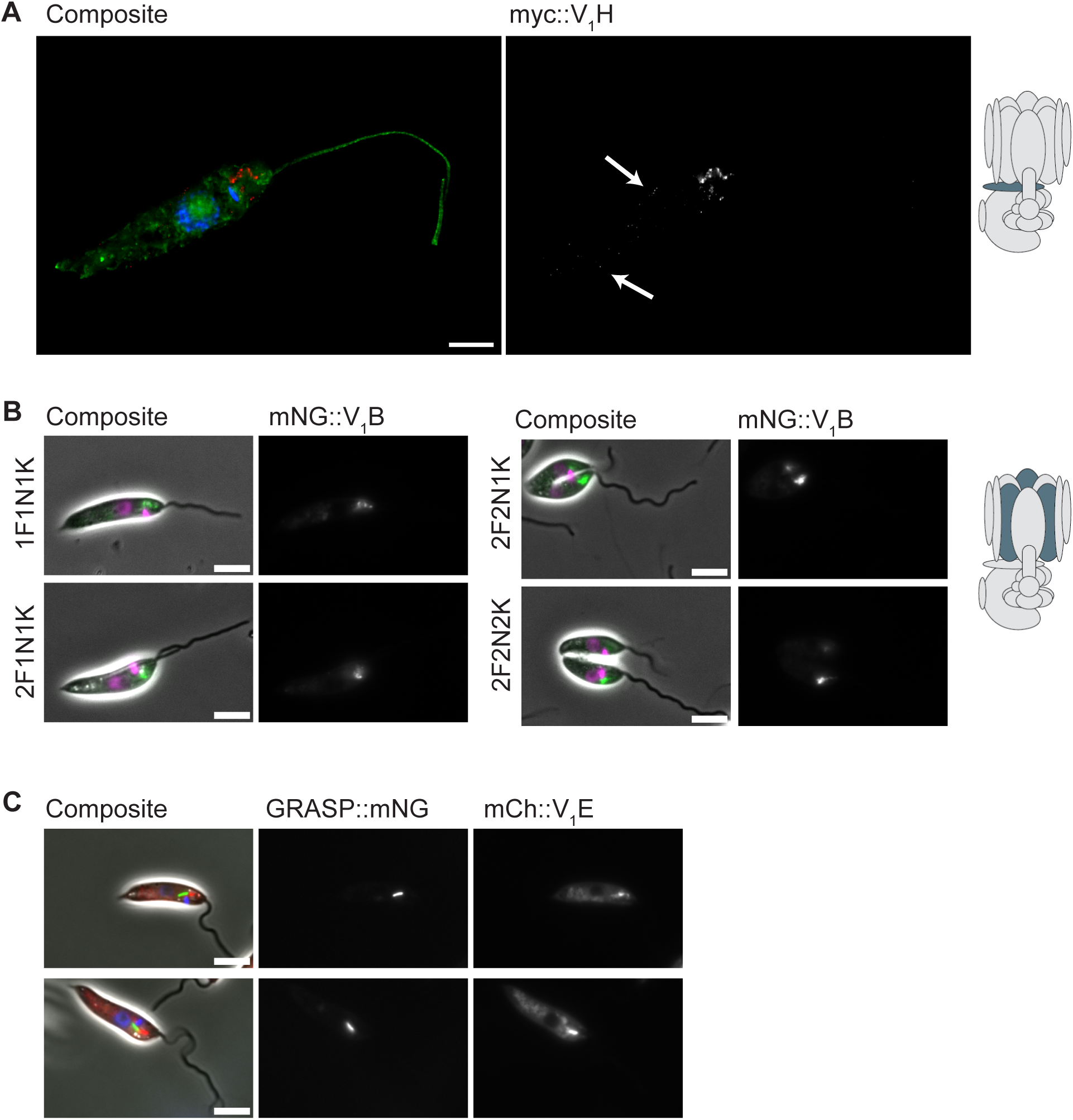
Subcellular localization of v-ATPase subunits. (**A**) Ultrastructure expansion microscopy showing localization of myc::V_1_H. Micrograph shows maximum intensity projection of z-stacks. The v-ATPase signal is in red. Arrows indicate the punctate signal in the cell body. Green: Atto-NHS-Ester stain, Blue: DNA. Scale bar is 10 µm. The cartoon highlights subunit H in the v-ATPase complex. (**B**) Promastigotes expressing mNG::V_1_B at different stages of the cell cycle, as determined by the number of flagella (F), nuclei (N) and Kinetoplasts (K). Magenta: DNA. Scale bars are 5 µm. The cartoon highlights subunit B. (**C**) Fluorescence micrographs of cells co-expressing mCherry-tagged V_1_E and mNG-tagged GRASP (Golgi) protein. Blue: DNA. Scale bars are 5 µm.

Analysis of the localisation pattern of mNG::V_1_B (LmxM.28.2430) during the cell cycle (Figure 2B) matches the description of CVC location and duplication during the *Leishmania* cell cycle as mapped in a 3D EM study (Hair et al., 2024). That study also defined the close proximity of the CVC to the Golgi apparatus, in the flagellar pocket region. We therefore hypothesized that the crescent-shaped structure where the v-ATPase signal is dominantly present in the cell is the CVC. We generated a cell line where GRASP (LmxM.32.2380) was tagged with mNG at the C-terminus as a Golgi apparatus marker (Halliday et al., 2019), and subunit V_1_E was tagged with mCh at the N-terminus. Fluorescence imaging of this double-tagged cell line indicated that the signals from both proteins are in proximity, but do not overlap (Figure 2C), in agreement with the location of the Golgi apparatus and CVC in the cell.

### v-ATPase partially colocalises with lysosomes, endosomes and a putative contractile vacuole protein

In *Leishmania*, no marker proteins for the CVC have been reported yet, however a proteome of the *Trypanosoma cruzi* CVC was reported by Ulrich et al (Ulrich et al., 2011). We endogenously tagged the *L. mexicana* homolog of one of the proteins found in the *T. cruzi* CVC, adaptor protein 180 (AP180, LmxM.24.0020), with mNG at the C-terminus. Colocalization analysis of AP180::mNG with mCh::V_1_E revealed a substantial overlap (median M2= 0.36, Figure 3A), suggesting that AP180 and the v-ATPase are in proximity, but possibly in different microdomains of the organelle, since the overlap between AP180 and V_1_E was not 100%.

**Figure 3:**
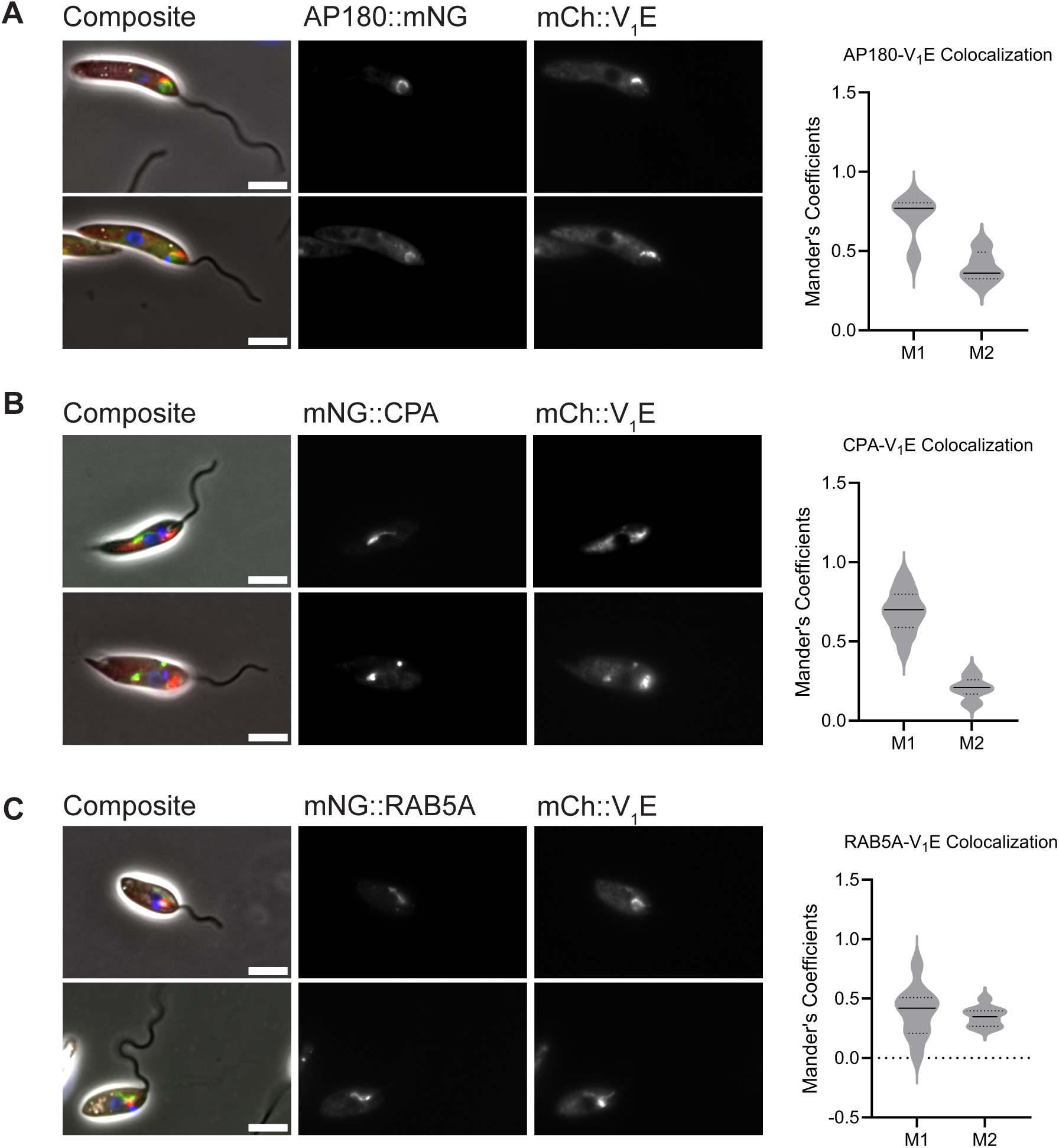
Colocalization of v-ATPase with organellar marker proteins. Fluorescence micrographs and colocalization analysis of mCh::V_1_E with (**A**) a putative contractile vacuole marker AP180, (**B**) lysosome marker CPA, (**C**) early endosome marker RAB5A. The graphs show measurements of signal overlap: Mander’s coefficient M1 shows the overlap of the signal from the green channel (organellar marker proteins) with the red channel signal (V_1_E). M2 is the overlap of red channel (V_1_E) with green channel signal (organellar marker proteins). Colocalization quantifications were done using Just another Colocalization Plugin (JaCOP) in ImageJ, on 5-15 different fields of view containing 7-15 cells. Median and quartiles are shown on the graphs. Blue: DNA. Scale bars are 5 µm.

Fluorescence microscopy of live cells and U-ExM showed a fraction of the v-ATPase signal in the cell body more distant from the FP region (Figure 2A), suggestive of localization to small cytoplasmic organelles. To map the v-ATPase to known organelles, we generated double-tagged cell lines co-expressing mCh::V_1_E and organellar marker proteins tagged with mNG (Figure 3, Supplementary figure 2, Supplementary table 1).

Analysis of a mNG::CPA (lysosome) mCh::V_1_E double-tagged cell line showed that a minor fraction of the V_1_E signal overlapped with the CPA signal (median M2= 0.21, Figure 3B). There were cell-to-cell variations in the level of colocalization between the CPA and V_1_E, which could indicate dynamic changes in v-ATPase occupancy and trafficking to and from the lysosome.

Since v-ATPases were found in endocytic vesicles and play essential roles in endocytosis in bloodstream form *T. brucei* (Xu et al., 2020), we next asked whether v-ATPase also localises to endosomes in *L. mexicana.* The V_1_E signal only partially overlapped with the signal for the early endosome marker RAB5A (LmxM.18.1130) (Halliday et al., 2019) (median M2= 0.35, Figure 3C). Similarly, a partial overlap was observed between V_1_E and RAB11 (LmxM.10.0910) (Supplementary Figure 2B).

As a flagellar pocket membrane marker, we tagged the clathrin heavy chain (LmxM.36.1630) alongside subunit V_1_E. There was partial overlap between the two proteins in some cells, however in most cases the two signals were distinct from each other, even though they were in proximity (Supplementary figure 2C).

Overall, these data suggest the *Leishmania* v-ATPase decorates multiple organelles, consistent with its canonically diverse functions. Some proportion of v-ATPases signal was found in the lysosomes and endosomes and the punctate localization pattern in the posterior part of the cell is compatible with an acidocalcisome localisation. Most of the v-ATPase signal was however in the flagellar pocket region, most likely within the CVC.

### v-ATPase knockout cells are arrested at the final stage of autophagy

Even though ΔV_1_E promastigotes were able to proliferate nearly at the same rate as parental controls in standard M199 medium (Figure 1), we noted a distinctive morphological phenotype: Live cell imaging of ΔV_1_E, V_1_E AB, and parental cells during their growth *in vitro* revealed the presence of large vacuole structures in the cell body of ΔV_1_E parasites starting from when the parasites reached a cell density of greater than 1x10^7^ cells/mL during their continuous growth in neutral or acidic pH (Figure 4A, B, supplementary figure 3A). Parental and V_1_E AB cells did not display such vacuoles on the same day of growth (Figure 4A, B, supplementary figure 3A). Interestingly, fewer than 10% of the ΔV_1_E cells had these structures while they were in a less dense cell culture environment on day 1 of growth in either neutral or acidic pH (Figure 4A, B, supplementary figure 3A). However, around 95% of the cells displayed these vacuoles when the cells reached stationary phase of growth, by day 3 (Figure 4B, supplementary figure 3A). In alkaline pH, these vacuolar structures were already present on day 1 in about 30% of the ΔV_1_E cells and then accumulated in more than 80% of the population on day 4 of growth (Figure 4B, supplementary figure 3B).

**Figure 4:**
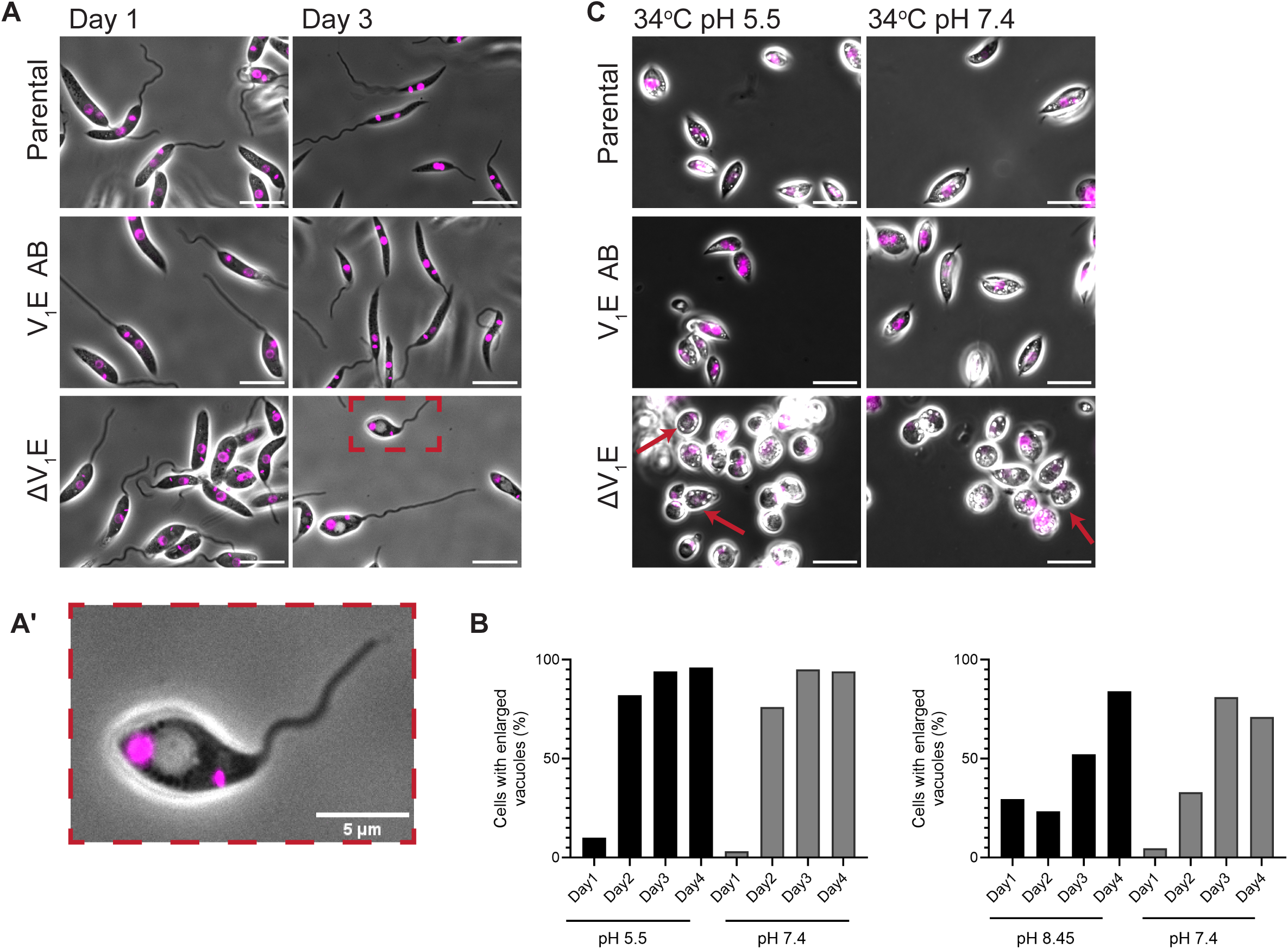
v-ATPase knockout leads to enlarged vacuolar structures. (**A**) Micrographs of parental, V_1_E AB and ΔV_1_E promastigotes grown in standard M199 medium at 27°C temperature for one (left) and three (right) continuous days. Scale bars are 10 µm. (**A’**) magnified view of a cell with large vacuole in the cell body. Magenta: DNA. (**B**) Quantification of parasites harboring enlarged vacuoles in pH 7.4 or pH 5.5 M199 medium over 4 days of growth. (**C**) Parental, V_1_E AB and ΔV_1_E parasites grown in pH 7.4 or pH 5.5 M199 medium at 34°C for 24 hours. Arrows indicate ΔV_1_E cells with enlarged vacuoles. Magenta: DNA. Scale bars are 10 µm.

In the ΔV_1_E parasites cultured at 34°C in neutral and acidic pH (Figure 4C), the vacuolar structures appeared already when the parasites were at cell densities less than 1x10^7^ cells/mL, in contrast to promastigotes kept at 27°C temperature (Figure 4A, C). Increasing the temperature to 34°C leads to changes in parasite morphology, where the cell body becomes rounded and the flagellum is shortened (Zilberstein and Shapira, 1994). Autophagy was shown to be required for differentiation to axenic amastigotes triggered by the combined exposure to acidic medium and elevated temperature (Williams et al., 2006). We thus hypothesized that these vacuoles indicate a defect in autophagy in ΔV_1_E cells that becomes apparent when cells are in stationary phase of growth, when the nutrients in the cell culture media are depleted, or when they are exposed to conditions that trigger cell remodelling and stage differentiation.

To investigate these vacuoles in more detail, we performed transmission electron microscopy (TEM) of the cells grown for 72 hours in neutral or acidic pH. TEM revealed the presence of partially degraded cellular material inside the enlarged vacuoles, such as fragments of the mitochondrion (Figure 5A, right panel, arrow). In addition, in some cases there were smaller vesicles that appeared to be fusing with the enlarged vacuolar structure, and a double membrane enclosing this enlarged vacuole (Supplementary figure 3C). All these observations are characteristic of autophagy-related structures, autophagosomes or autolysosomes.

**Figure 5:**
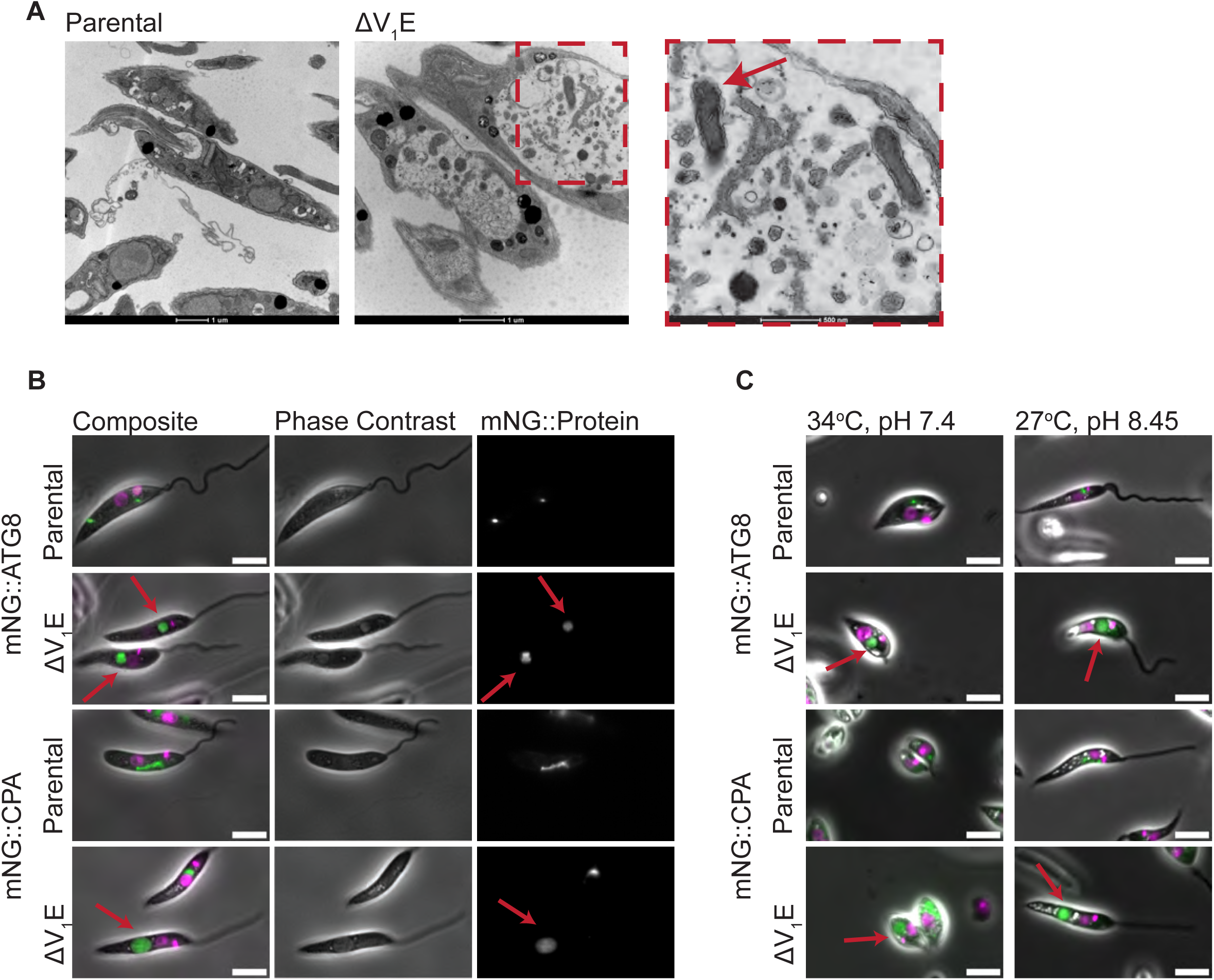
The enlarged vacuolar structures of the v-ATPase knockouts are autolysosomes. (**A**) Transmission electron microscopy micrographs of parental and ΔV_1_E promastigotes grown for three continuous days in standard M199 medium at 27°C. Arrow shows a mitochondrion piece within the vacuole. (**B**) Fluorescence micrographs showing parental or ΔV_1_E promastigotes expressing mNG::ATG8 (autophagosome marker) or mNG::CPA (lysosome marker), after three continuous days in standard M199 medium at 27°C. (**C**) Tagged cells as in (B) grown for one day in standard M199 medium at 34°C (left), or for three continuous days in M199 medium at pH 8.45, 27°C (right). Arrows indicate the enlarged vacuoles. Magenta: DNA. Scale bars are 5 µm.

We studied these structures further by tagging autophagosome marker ATG8 (LmxM.19.1630) endogenously (Williams et al., 2006), or a marker of the MVT lysosome CPA (LmxM.19.1420) (Halliday et al., 2019), with mNG. We then deleted V_1_E in these autophagy marker cell lines. In neutral pH medium, the enlarged vacuoles that formed when cells reached the stationary phase of growth were positive for both mNG::ATG8 and mNG::CPA signal (Figure 5B). Similarly, the vacuoles observed in 27°C, pH 8.45 in stationary phase cultures and at 34°C, pH 7.4 on day 1 of growth were also positive for mNG::ATG8 and mNG::CPA signal (Figure 5C).This indicates that these are autolysosomes, the final structures that are formed in the autophagy pathway before lysosomal degradation.

To further confirm that the enlarged vacuoles in ΔV_1_E cells correspond to autolysosomal structures, we generated an autophagy-deficient mutant, by deletion of ATG5 (LmxM.29.0980). ATG5 functions in the ATG8 lipidation pathway, where it forms a complex with ATG12, and together they act as the E3-like ligase for conjugation of ATG8 to phosphatidylethanolamine (Hanada et al., 2007). In *Leishmania major*, ATG5 was shown to be essential for formation of ATG8 puncta, indicating its role in autophagosome formation (Williams et al., 2012). We generated ΔATG5 cell line in mNG::ATG8 expressing parentals, as well as in C9T7, and observed that mNG::ATG8 puncta disappeared upon ATG5 knockout, as has been previously reported (Williams et al., 2012) (Supplementary figure 3D). Moreover, treatment of the cells with BafA1 for 3 days, which phenocopies the v-ATPase knockout phenotype leading to formation of ATG8-labelled enlarged vacuoles, showed that these compartments did not form in the ΔATG5 cell lines (Supplementary figure 3D). This indicates that the formation of enlarged vacuoles resulting from v-ATPase disruption or inhibition requires a functional autophagosome biogenesis pathway.

Taken together, this suggests that *Leishmania* lacking functional v-ATPase become arrested at the final stage of autophagy, where autolysosomes can form but get stalled, most likely due to dysfunction in lysosome enzyme activity. We hypothesized that this was due to an increase in the luminal pH of the lysosome in ΔV_1_E cells.

### L. mexicana lysosomes have an average pH of 5.6

The MVT lysosome of *Leishmania* differs from the mammalian lysosomes in morphological aspects. Vesicles constituting the lysosome assemble to form the elongated, tubular shape of the MVT lysosome, a structure that is dynamic throughout the cell cycle (Mullin et al., 2001; Wang et al., 2020b). Even though the morphology and dynamics of the MVT lysosome have been studied in detail (Wang et al., 2020b), the pH of this organelle in *Leishmania* remained unknown. It has been suggested that the MVT has a luminal pH higher than 5.5, due to the observations that lysotracker dyes, which stain subcellular compartments of pH 5.5 and below, failed to stain the lysosome in *Leishmania* (Mullin et al., 2001). To measure the lysosome pH in *L. mexicana*, we utilized a ratiometric, genetically encoded pH biosensor called pHLuorin2 (pHL2), a GFP-derivative fluorophore, whose light excitation spectrum is pH-dependent (Mahon, 2011). We endogenously tagged CPA with pHL2 to target the fluorophore to the lysosome and confirmed that the biosensor was fluorescent in the lysosome when excited with 395 nm or 475 nm (Figure 6A). pH measurement experiments using pHL2::CPA cell line showed a mean pH of 5.6 in the lysosome (Figure 6B). The pH of individual lysosomes ranged from pH 4.0 to pH 6.5, with the majority of lysosomes in the range of pH 5.6-6.2 (Figure 6C). To investigate possible underlying reasons for the variations in the lysosome pH we first compared lysosomes with different morphology, tubular or vesicular. There was no significant difference in the pH between the two populations (Figure 6D, left panel). Next, we compared the lysosome pH in different cell cycle stages, comparing cells with 1 flagellum vs 2 flagella. Again, there was no significant difference (Figure 6D, right panel). Currently the underlying mechanism of the pH heterogeneity between individual lysosomes remains unclear.

**Figure 6:**
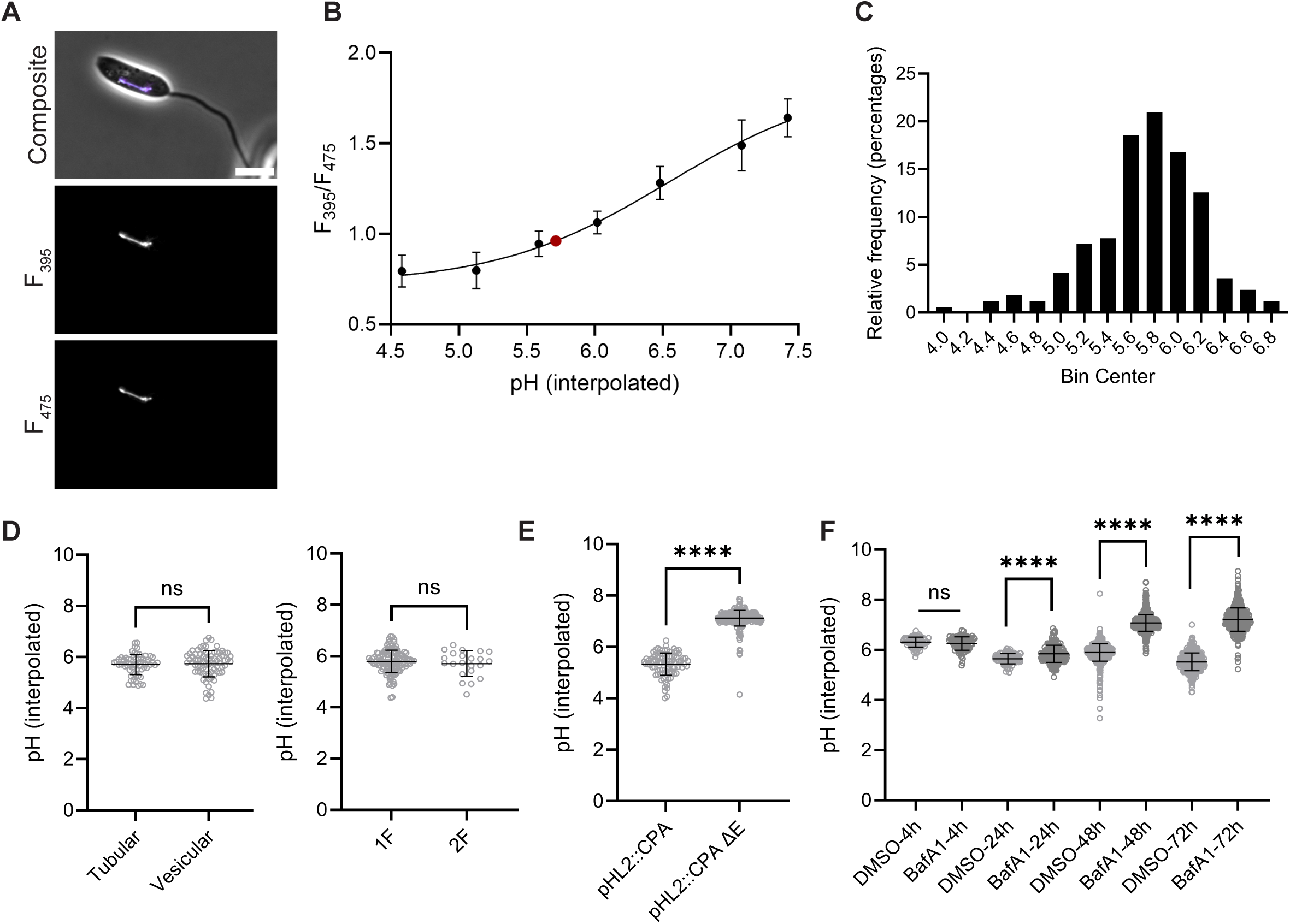
Perturbation of v-ATPase function leads to an increase in lysosomal pH. (**A**) Fluorescence micrographs showing localization of pHLuorin2 (pHL2)::CPA at the MVT lysosome. Scale bar is 5 µm. (**B**) Calibration curve and pH measurement of pHL2::CPA in *L. mexicana*. Each point represents the mean F_395_/F_475_ signal intensity ratios from n=40-180 cells. Error bars indicate the standard deviation. The red dot indicates the interpolated lysosomal pH of pHL2::CPA samples imaged in PBS. The graph is representative of four independent calibration curves and pH measurements. (**C**) Histogram of individual pH measurements from (B). (**D**) Distribution of individual lysosome pH measurements according to lysosome morphology (left) and cell cycle stage, as assessed by the presence of 1 or 2 flagella (F) (right). (B) Lysosome pH measurements of pHL2::CPA in parental cells and in the ΔV_1_Emutant. (**F**) Lysosome pH measurements of cells treated with BafA1 or DMSO (control) for 4 hours, 24 hours, 48 hours and 72 hours.

### Maintenance of lysosomal pH requires a functional v-ATPase complex in L. mexicana

To investigate whether lysosomal pH depends on the function of the v-ATPase, we employed two complementary approaches. First, we generated a V_1_E knockout in pHL2::CPA expressing cells and measured the lysosome pH. In the pHL2::CPA ΔV_1_E cells, the mean lysosome pH increased to 7.1 (Figure 6E), indicating that v-ATPase plays a crucial role in acidifying the lysosome pH.

Next, we treated pHL2::CPA expressing cells with BafA1. We measured the lysosome pH at time points 4, 24, 48 and 72 hours of drug treatment. There was no significant change in pH after 4 hours of BafA1 treatment, indicating that the drug did not have an acute effect on lysosome pH (Figure 6F). By 24 hours, however, the lysosome pH became significantly higher (mean pH =7.0), an effect that was maintained up to 72h (Figure 6F). BafA1 treated cells also displayed the enlarged vacuoles, similar to ΔV_1_E cells (Supplementary figure 3E). This shows that pharmacological inhibition of v-ATPase by BafA1 or deletion of V_1_E both result in a neutralized lysosome pH and similar morphological changes.

### v-ATPase knockout phenotype is phenocopied by Bafilomycin A1 in L. mexicana and Trypanosoma brucei

We next asked whether BafA1 treatment in *Leishmania* prevents fusion of autophagosomes and lysosomes. We treated mNG::CPA mCh::ATG8 double-tagged promastigotes with BafA1. Live cell imaging of this cell line showed that mNG::CPA and mCh::ATG8 signals overlapped in the BafA1 treatment group, indicating that the drug did not prevent autophagy initiation and fusion of autophagosomes with lysosomes (Figure 7A).

**Figure 7:**
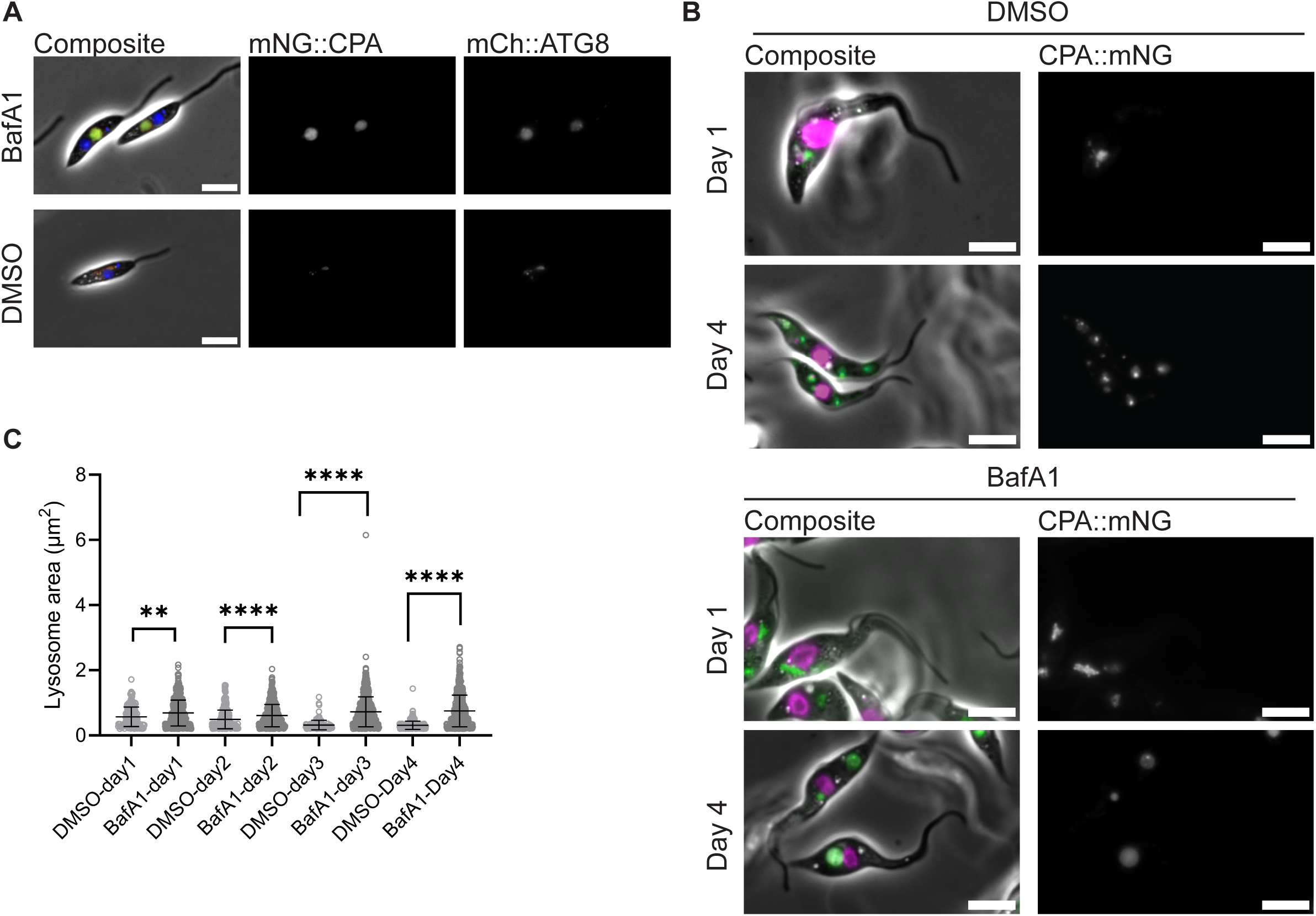
BafA1 does not prevent autophagosome-lysosome fusion in *L. mexicana* and phenocopies v-ATPase knockout phenotype in *T. brucei*. (**A**) Fluorescence micrographs of *L. mexicana* promastigotes co-expressing mNG::CPA (lysosome marker) and mCh::ATG8 (autophagosome marker) tags, treated with BafA1 or DMSO (control) for three continuous days (**B**) Fluorescence micrographs showing procyclic form *T. brucei* expressing CPA::mNG, treated with DMSO (control) or 160 nM BafA1 for 1 day and 4 continuous days. Magenta: DNA. Scale bars are 5 µm. (**C**) Quantification of *T. brucei* mNG::CPA area (lysosomes) from (B). Magenta: DNA. Scale bars are 5 µm.

In procyclic form (PCF) *Trypanosoma brucei*, another kinetoplastid, epitope tagging and colocalization analysis of v-ATPase subunit a (V_o_ domain) showed partial colocalizations with acidocalcisomes, Golgi apparatus and lysosomes (Huang et al., 2014). To test whether the inhibition of v-ATPase function by BafA1 treatment would also lead to enlargement of lysosomes in *T. brucei* PCF, we endogenously tagged *T. brucei* CPA (Tb927.6.100) on the C-terminus with mNG, as a lysosome marker (Billington et al., 2023). Treatment of the CPA::mNG expressing parasites with 160 nM BafA1 or DMSO as a control for 24, 48, 72 and 96 hours led to an increase in lysosome size increased over time from day 1 to day 4 (Figure 7B,C).

Taken together these results suggest a conserved essential function for trypanosomatid v-ATPases in lysosomal acidification, which is required for the resolution of autolysosomes.

## Discussion

Recent work showed that the v-ATPase is essential for the survival of amastigote form *L. mexicana* in iPSC-derived macrophages, mice (Albuquerque-Wendt et al., 2025) and in *Lu. longipalpis* sandflies (Sadlova et al., 2026). Moreover, ΔV_1_E parasites failed to differentiate into metacyclic form promastigotes *in vivo*, based on morphological analysis of the parasites isolated from sandflies; and they also did not express the metacyclic stage marker gene SHERP in stationary phase culture *in vitro* (Sadlova et al., 2026).

Since differentiation is often triggered by environmental stress, and we hypothesized that the v-ATPase functions in cellular stress responses, we exposed the v-ATPase knockout mutants to adverse conditions of the kind they encounter on their journey from arthropod gut to mammalian phagosome. The growth phenotype of ΔV_1_E mutants in acidic pH or alkaline pH at 27°C showed that the v-ATPase complex is important in conditions where the external pH deviated from neutral; the ectopic re-expression of the deleted gene rescued this phenotype. In addition, we showed that ΔV_1_E mutants could not adapt to heat (34°C temperature) in neutral or acidic pH. Moreover, prolonged culture in stationary phase and exposure to hyperosmotic medium were detrimental to v-ATPase mutants. This indicates that ΔV_1_E promastigotes fail to tolerate diverse stress conditions, including those that lead to *in vitro* differentiation of metacyclics and axenic amastigotes. This is in line with our previous findings that ΔV_1_E mutants are deficient in metacyclogenesis *in vitro* and *in vivo* (Sadlova et al., 2026) and explains why v-ATPase function is essential in the parasites’ varied natural environments outside well-buffered temperature-controlled laboratory cultures.

The diverse nature of the stresses that reveal a need for v-ATPase function suggest either that the v-ATPase has independent functions in different pathways or that it is crucial for a mechanism downstream of disparate stressors. Autophagy initiation is such a point of convergence, with a well-documented function for v-ATPases in other species. While early events in autophagy have been described, and their importance for parasite differentiation, stress response and infectivity studied in different *Leishmania* species (Giri and Shaha, 2019; Williams et al., 2013; Williams et al., 2012), the role of the *Leishmania* v-ATPase in lysosomal degradation and autophagy was assumed but remained poorly defined. Our results provide direct evidence from genetic mutants that the v-ATPase is involved in the autophagy pathway in *L. mexicana:* We observed by live cell imaging and TEM, that ΔV_1_E mutants accumulated enlarged vacuolar structures in the cell body in response to being in dense *in vitro* cultures, hyperosmotic stress, or heat. The formation of these vacuoles was dependent on ATG5 and they were positive for the autophagosome marker ATG8 and lysosome marker CPA, indicating that autolysosomes can fuse with the lysosome in ΔV_1_E parasites, and expand, but autolysosomes cannot be resolved.

Measurements of the luminal pH of the lysosomes showed an increase from 5.6 to 6.6-7.2 when the v-ATPase function was perturbed by genetic deletion or pharmacological inhibition. A plausible mechanistic explanation for the phenotype is that the neutralisation of the lysosomal pH renders lysosomal digestive enzymes unfunctional. Similar large “apoptotic bodies” were first described in protease deficient yeast (Takeshige et al., 1992) and were also seen in *L. mexicana* upon inhibition of cysteine peptidases (Williams et al., 2006). Another contributing factor could be impaired export of the material out of the lysosome, which may also be pH-dependent. The efflux pumps that mediate recycling of lysosomal content to the cytoplasm are still poorly characterized in most species. The amino acid efflux pump Atg22 (Yang et al., 2006) is restricted to yeast, while in mammals the v-ATPase is required for an mTORC1-dependent mechanism of coupling amino acid sensing to efflux from lysosomes via amino acid transporter SLC38A9 (Wyant et al., 2017). Which of the many uncharacterised amino acid transporters (Albuquerque-Wendt et al., 2025) fulfil similar roles in *Leishmania* remains to be studied.

Our results show that the effect of v-ATPase inhibition on lysosome pH regulation and autophagy in *L. mexicana* is profound. It appears however that the primary subcellular location of the v-ATPase is not the lysosome, but rather the flagellar pocket region, as shown by endogenous tagging and live cell imaging experiments for subunits V_1_G and V_1_H in promastigotes and axenic amastigotes (Albuquerque-Wendt et al., 2025). Here we show the same localization for seven of the eight V_1_ subunit proteins and for four of nine predicted V_o_ proteins. The typically diffuse and weak fluorescence seen for the remaining subunits could be an artefact of tagging, particularly in membrane-associated V_o_ domain proteins.

Taken together, the available evidence suggests that the v-ATPase complex proteins in the vicinity of the FP localise to the CVC. Colocalization analysis of V_1_E with the putative CVC protein AP180 showed overlap, albeit not a 100% colocalisaiton. This could be explained by the proteins localising to the different microdomains of the CVC, or by the dynamic nature of this organelle. Further study of the *Leishmania* CVC is warranted to better understand its functions in relation to other cell organelles and contribution to *Leishmania* cell physiology. In this study we found that ΔV_1_H mutant promastigotes are more sensitive to hyperosmotic stress, compared to parental controls. Given the conserved role of the contractile vacuoles in osmoregulation in many protists (More et al., 2024), and the findings that its membrane is decorated by v-ATPase molecules in other protists (Docampo et al., 2013; Fok et al., 1995; Plattner, 2013), it seems plausible that v-ATPase complexes localised on the CVC membrane in *Leishmania* may function in osmoregulation.

v-ATPases are also known to decorate the membranes of acidocalcisomes in *T. brucei* and *T. cruzi* (Docampo et al., 1995; Huang et al., 2014; Vercesi et al., 1994). Acidocalcisomes, acidic organelles that serve as inorganic cation and phosphate storage compartments, have also been shown to be involved in osmoregulation in *T. cruzi* (Li et al., 2011; Rohloff et al., 2004). They fuse with the contractile vacuole in response to hypoosmotic stress, thereby contributing to the export of excess water via the contractile vacuole (Rohloff et al., 2004). It is likely that v-ATPase complexes decorating the acidocalcisome membrane contributed to the more dispersed signal in the posterior part of the cell body, which we observed by live cell imaging and expansion microscopy. The involvement of acidocalcisomes in osmoregulation in *Leishmania* and the specific contributions of the v-ATPase to acidocalcisome physiology and function should be explored in the future.

In light of the importance of the v-ATPase in the acidification of the lysosome, it might seem contradictory at first that only a minor fraction of v-ATPase seems to localise at the lysosome. It should be noted, however, that the average lysosome pH in *L. mexicana* is less acidic compared to mammalian lysosomes, suggesting that a small number of v-ATPase complexes at the lysosome membrane might be sufficient to regulate its pH. In addition, we have also shown that a fraction of the v-ATPase pool colocalises with RAB5A and RAB11, suggesting endosomal localisation of the protein complex. Therefore, it could be indirectly contributing to the acidity of the lysosome, by acidification of endocytic vesicles, which eventually fuse with the lysosome. The CVC, acidocalcisomes and the MVT lysosome of *Leishmania* are dynamic organelles connected through a network of vesicular trafficking routes whose dynamic regulation in response to changing environmental signals is not fully known in *Leishmania*. The CVC could act as a “hub” of v-ATPase storage, and the v-ATPase molecules might be trafficked to their destinations such as the lysosome or endosomes upon need. To discover whether the v-ATPase is a driver or passenger, and the mechanisms controlling v-ATPase activity in trypanosomatids, will require studies of these dynamic processes in living cells.

Taken together, our data indicates that v-ATPase serves essential functions in *Leishmania* cell biology. Without a functional v-ATPase complex, parasites are unable to withstand stress conditions and complete their life cycle. The major contributor to this phenotype is very likely the deficiency in the autophagy pathway. Autophagy is a general pro-survival stress response, and due to their defect in this pathway, v-ATPase knockout mutants are sensitive to stress conditions such as hyperosmotic stress and heat. The importance of v-ATPase in *Leishmania* fitness and physiology, however, is likely to be more than only its role in autophagy. v-ATPases contribute to many other essential cellular processes such as endocytosis and protein sorting across eukaryotes, from *T. brucei* to yeast and human cells (Currier et al., 2018; Forgac, 2007; Lecordier et al., 2014; Xu et al., 2020) and likely in *Leishmania* too.

Finally, given the essentiality of v-ATPase in parasites’ adaptation to stress conditions, completion of the life cycle and the role in autophagy, this work highlights the v-ATPase as a potential drug target. Identification of targetable kinetoplastid- or *Leishmania*-specific v-ATPase subunit features, associated factors or proteins that regulate its function would be necessary to specifically target the *Leishmania* v-ATPase without interfering with the host v-ATPase.

## Materials and Methods

### *Cell culture of* Leishmania mexicana

*L. mexicana* promastigote form Cas9T7 cell line (parental), and the generated tagged or knockout mutants were grown in T25 cm^2^ flasks at 27°C or in 24-well plates at 27.5°C, 5%CO_2_ incubators, in filter-sterilized M199 medium (Gibco) supplemented with 10% fetal bovine serum (FBS), 2.2 g/L NaHCO_3_, 0.005% hemin and 40 mM 4-(2-Hydroxyethyl)piperazine-1-ethanesulfonic acid (HEPES) pH 7.4 (standard M199 medium). The parasites were maintained at a density of 1x10^6^-1x10^7^ cells/mL by regular passages. Axenic amastigote cultures were generated by splitting promastigotes in complete M199 medium pH adjusted to 5.5 by HCl, at 34°C + 5% CO_2_.

### *Cell culture of procyclic form* Trypanosoma brucei

*T. brucei* Lister 427 29-13 (Wirtz et al., 1999) were maintained in filter-sterilized SDM79 medium (Bioconcept/Amimed 9.04V01-M) supplemented with 10% FBS and 0.001 mg/ml hemin, pH adjusted to 7.3. The cells were grown in T25 cm^2^ flasks at 27°C or in 24-well plates at 27.5°C, 5% CO_2_ incubators.

### Cloning of pHLuorin2 into pPLOT plasmid

Plasmid pPLOT-pHLuorin2 was generated by replacing the mCherry gene in pPLOT-mCherry-Puromycin with the pHLuorin2 coding sequence by restriction digestion cloning using the *Hind*III and *Bam*HI sites. The pHLuorin2 sequence was amplified from plasmid pLEW-pHluorin2-PTS1 (a gift from Kenneth Christensen, Addgene plasmid #213776, http://n2t.net/addgene:213776 ; RRID:Addgene_213776) and restriction enzyme cut sites for *Hind*III and *Bam*HI were inserted at the 5’ and 3’ end of the pHLuorin2 coding sequence, respectively, with the following primers:

pHL2_Forw_*Hind*III 5’-TCAGTGAAGCTTATGAGCAAAGGCGAAGAATTGTTTAC-3’

pHL2_Rev_*Bam*HI 5’-CAGTCTGGATCCTTTGTAAAGTTCGTCCATGCC-3’

### *Generation of endogenously tagged and deletion mutants by CRISPR-Cas9 & diagnostic PCR for knockout verification in* L. mexicana

Gene deletions and endogenous tagging were performed as described in (Beneke and Gluenz, 2019). The sequences of primers for generation of donor DNA and 5’ or 3’ sgRNA template were retrieved from LeishGEdit (http://www.leishgedit.net). For endogenous tagging, donor DNAs were amplified from pPLOTv1 blast-mNeonGreen-blast, pPLOTv1 puro-mCherry-puro or pPLOT puro-pHLuorin2-puro plasmids. For knockouts, pTBlast_v1, pTPurov1 and/or pTNeov1 plasmids were used as templates for amplification of donor DNAs. Successful transfectants were selected by addition of corresponding antibiotics Puromycin (final concentration 20 μg/ml), Blasticidin (final concentration 5 μg/ml) and/or G418 (final concentration 40 μg/ml) 16 hours post transfection, and cells were kept in the presence of drugs for at least three passages. For confirmation of knockout cell lines, ORF verification primers were designed for the amplification of 200-300 bp region of the gene of interest. ORF verification primers used in this study can also be found in the supplementary table 2, and ORF verification results for newly generated knockout lines are in supplementary figures 4 and 5. For the diagnostic PCRs, master mixes were prepared using GoTaq G2 Hot Start Green Master Mix (Promega), according to the manufacturer’s protocol. As a technical positive control for the diagnostic PCRs, a PCR reaction to amplify the IFT88 gene (Gene ID: LmxM.27.1130) was set up.

### *Endogenous tagging of PCF* T. brucei

For the endogenous tagging of CPA (Tb927.6.100), tagging construct was generated using pPOTv7 encoding Puromycin drug resistance marker and mNeonGreen. Primer sequences for the amplification of the tagging construct were retrieved from TrypTag.org (Billington et al., 2023). TbPCF 29-13 parental cell line was transfected with 10 µg PCR construct, in a transfection buffer mix (90 mM NaPO_4_ (pH 7.4), 50 mM HEPES (pH 7.4), 5 mM KCl, 0.15 mM CaCl_2_), using Amaxa Nucleofector electroporation system, with Nucleofector X-014 program. Successful transfectants were selected by addition of puromycin at a final concentration of 1 µg/mL 24 hours post-transfection. The cells were cloned by limiting serial dilution on a 96 well-plate to obtain clonal cell populations. For cloning, fresh SDM79 medium mixed with an SDM79 medium isolated from a stationary phase culture of *T. brucei* 29-13 PCF in a 1:5 ratio was used.

### *Growth curves* in vitro

Parental, ΔV_1_E and V_1_E AB cell lines were seeded in 3 identical flasks (triplicates) at a density of 1x10^5^ or 5x10^5^ on day 0 in complete M199 medium. Where needed, M199 medium pH was adjusted to 5.5 by addition of HCl or to 8.45 by addition of NaOH. Parasites were grown at 27°C or 34°C + 5% CO_2_ conditions. Every 24 hours parasite growth was assessed by counting the cells with CASY^®^ cell counter (Cambridge Bioscience) using a 60 μm capillary and measurement range set between 2 and 15 μm, for indicated time points. Where needed, parasite cultures were diluted back to their initial density and parasite growth was assessed as described above for the indicated time points.

### Fluorescence microscopy and colocalization analysis

Live cell imaging was performed as described in (Albuquerque-Wendt et al., 2025). Cells were imaged either on an inverted Nikon Eclipse Ti2-microscope equipped with Kinetix camera and 100X oil objective, numerical aperture (NA) 1.3, or with Leica DM6000 B microscope equipped with Leica DFC360 FX camera and 100X oil objective, NA 1.3. Micrographs were analyzed with ImageJ software. For the colocalization analysis, JaCOP (Just another Colocalization Plugin) was used. Mander’s coefficients M1 and M2 were plotted in GraphPad Prism.

### Lysosome pH measurements by fluorescence microscopy

The pH of lysosomes was measured with the “ratiometric” pH biosensor pHLuorin2. Acidification of the environment surrounding the fluorophore pHL2 leads to a decrease in excitation in 395 nm light, with a subsequent increase in excitation in 475 nm; both excitation wavelengths result in light emission at ∼510 nm (Mahon, 2011). *L. mexicana* promastigotes or axenic amastigotes expressing pHLuorin2::CPA were harvested by centrifugation at 800 *g* for 5 minutes. The cells were washed twice with PBS and resuspended in universal pH calibration buffers (15 mM 2-(N-Morpholino)ethanesulfonic acid (MES), 15 mM HEPES, 130 mM KCl) with pH values ranging from pH 4.0 to 7.5, supplemented with 10 μM nigericin and 4 μM valinomycin (for the calibration curve), or in PBS and incubated 5 minutes at 27°C before live cell imaging (Call et al., 2024). Where indicated, pHluorin2::CPA expressing parasites were treated with 400 nM BafA1 for 24h, 48h or 72h; or pHluorin2::CPA ΔV_1_E cells were used for live cell imaging and subsequent pH interpolations. The cells were seeded in glass bottom 24-well plates for live cell imaging. For illumination of the pHLuorin2 fluorophore, light excitation and emission parameters were set as the following: GFP channel (excitation: 475 nm emission: 510 nm), DAPIex_GFPem channel (excitation: 395 nm emission: 510 nm). Image acquisitions were automated with Nikon JOBS software. Images were analyzed with ImageJ software (Schindelin et al., 2012), using a macro to segment the pHLuorin2::CPA signal and measure signal intensities of the relevant channels. Mean signal intensity ratios of DAPIex_GFPem/GFP channels were plotted against the known pH values, and the unknown pH was calculated by interpolation of the signal intensity ratio in GraphPad Prism as previously described (Call et al., 2024).

### Ultrastructure expansion microscopy

Ultrastructure expansion microscopy (U-ExM) protocol was adapted from (Gambarotto et al., 2021) and was performed as described in (Fochler et al., 2026), with the following changes: Antibody staining was performed before gelation of the sample. Primary antibody staining was done with mouse anti-c myc 4A6 monoclonal antibody (Sigma Millipore, Cat. No: 05-724-25UG) in 2% BSA in PBS blocking solution for 1 hours at room temperature, followed by 3 times 5 minutes washes with PBS-T. Secondary antibody staining was performed with goat anti-mouse IgG AlexaFluor 594 antibody (Invitrogen, Cat. No: A11005) for 1 hour at room temperature in dark. NHS-Ester (final concentration 1 µM, Atto 488 NHS Ester Cat. No: AD488 31) and DNA staining (final concentration 20 µg/ml, Hoechst 33342) were done after gelation of the sample, at 37°C temperature for 1.5 hours.

### MTT assay

Parasite viability in hyperosmotic stress was measured with an MTT (3-(4,5-dimethylthiazol-2-yl)-2,5-diphenyl-2H-tetrazolium bromide) assay. Briefly, M199 cell culture media osmolality was adjusted to 1600 mOsm by addition of sucrose in the media (hyperosmotic stock M199), and this hyperosmotic stock M199 was diluted with standard M199 medium (osmolality 340 mOsm) via serial dilutions in a 96-well plate. *L. mexicana* C9T7 and ΔV_1_H promastigotes in standard M199 were seeded in the 96-well plate containing serial dilutions of hyperosmotic M199 at a final seeding density of 1 x 10^6^ cells/mL, in six technical replicates. As a positive control, the cells were seeded in standard M199 medium. To account for the blank absorbance coming from sucrose-added M199 medium, a blank 96-well plate containing no cells but M199 medium with the corresponding osmolalities were prepared. The cells were incubated in the 96-well plates for 24 hours at 27°C + 5% CO_2_ before addition of the MTT reagent. After adding MTT at a final concentration of 0.650 mg/mL, the cells were incubated for another 3 hours at 27°C + 5% CO_2_ and all the subsequent steps were performed in dark. The reaction was stopped by addition of SDS at a final concentration of 3.5% (w/v). The absorbance was measured at 570 nm using a plate reader. For each osmolality condition, the average absorbance from the blank control was subtracted for background correction. Parasite viabilities were plotted with GraphPad Prism, where the values were normalized according to untreated positive control (100% viability).

### Transmission electron microscopy (TEM) sample preparation and image acquisition

*Leishmania* promastigotes were fixed in 2.5% glutaraldehyde, 2% formaldehyde in 100 mM phosphate buffer pH 7.4 as described by Hoog et al. (2010). The fixed pellets were washed in 0.1 M cacodilate buffer, and post-fixed in 1% osmium tetroxide (OsO_4_) and 1.25% potassium ferrocyanide (vol:vol) in the same buffer for 1 hour in the dark, then washed with distilled water and contrasted *en bloc* with 0.5% aqueous uranyl acetate for 1 hour at room temperature in the dark. The samples were then dehydrated in a series of increasing acetone concentrations (30%, 50%, 70%, 90% and four times in 100%, incubating for 15 minutes in each step), and embedded in epoxy resin. Ultra-thin sections (70 nm) were cut on a Leica Ultramicrotome, collected on formvar-coated mesh grids and contrasted with 2% aqueous uranyl acetate. Electron micrographs were acquired on a FEI Tecnai Spirit BioTwin electron microscope, using either EAGLE or VELETA camera and analysed with ImageJ software (Schindelin et al., 2012).

## Acknowledgements

Thanks to Jeremy Mottram for helpful discussions, Torsten Ochsenreiter (University of Bern) for providing the *T. brucei* PCF 29-13 cell line and Leoandro Lemgruber (Glasgow Imaging Facility, University of Glasgow), and Beat Haenni (Univeristy of Bern) for help with TEM sample preparation. Microscopy was performed on equipment supported by the Microscopy Imaging Center (MIC), University of Bern, Switzerland.

## Supplementary Data

**Supplementary Figure 1: Subcellular localization of v-ATPase subunits**

**Supplementary Figure 2: Partial colocalisation of v-ATPase with organellar marker proteins**

**Supplementary Figure 3: Enlarged vacuolar structures of v-ATPase knockouts are observed under different conditions and are identified as autolysosomes**

**Supplementary Figure 4: Diagnostic PCR for V_1_E knockout verification**

**Supplementary Figure 5: Diagnostic PCR amplification of ATG5 for knockout verification Supplementary Table 1: Localisation summary**

**Supplementary Table 2: KO cell lines and primers**

