## Supplementary Figures 1-5 for "Vacuolar type H^+^ ATPase is involved in stress responses in *Leishmania mexicana* by regulating the lysosomal pH"

### Supplementary Data

#### Supplementary Figure 1: Subcellular localization of v-ATPase subunits

Fluorescence micrographs of *L. mexicana* promastigotes expressing v-ATPase subunits of the $V_1$ domain tagged with mNeonGreen (mNG) or mCherry (mCh) (**A**) at the C- terminus or (**B**) at the N-terminus. (**C**) The parasites expressing C-terminally tagged $V_1$ domain subunits imaged after growth in axenic amastigote differentiation conditions (34°C, pH 5.5) for 72 hours prior to imaging. (**D**) Axenically differentiated parasites with $V_1$ domain subunits tagged at the N- terminus. (**E**) Fluorescence micrographs of promastigotes expressing v-ATPase subunits of the $V_o$ domain tagged with mNG at the C- or N- terminus. (**F**) Axenically differentiated parasites with tagged $V_o$ domain subunits. Scalebars = 5 µm.

#### Supplementary Figure 2: Partial colocalisation of v-ATPase with organellar marker proteins

(**A**) Fluorescence micrographs of *L. mexicana* promastigotes expressing v-ATPase subunit $V_od$ tagged with mNG at the C- terminus and subunit $V_1E$ tagged with mCh at the N- terminus show overlap of the two signals. (**B**) Promastigotes co-expressing RAB11 (Gene ID: LmxM.10.0910) tagged with mNG at the C- terminus and v-ATPase subunit $V_1E$ tagged with mCh at the N- terminus show partial overlap of the two signals. (**C**) Promastigotes expressing clathrin heavy chain (Gene ID: LmxM.36.1630, flagellar pocket marker) tagged with mNG at the C- terminus and $V_1E$ tagged with mCh at the N- terminus.

#### Supplementary Figure 3: Enlarged vacuolar structures of v-ATPase knockouts are observed under different conditions and are identified as autolysosomes

(**A**) Micrographs of parental, $V_1E$ add-back (AB) and $\Delta V_1E$ promastigotes grown at 27°C for 1 day and 3 days in pH 5.5 medium or (**B**) in pH 8.45 medium. Large vacuoles are present in $\Delta V_1E$ cells. Magenta: DNA. Scale bars: 10 µm. (**C**) TEM micrographs of vacuoles in $\Delta V_1E$ promastigotes grown for 3 days in the indicated pH conditions. Arrows show small vesicular structures apparently fusing with the large vacuole. (**D**) Top, micrographs of parental and autophagy-deficient ΔATG5 promastigotes exposed for 3 days to BafA1 treatment, or DMSO as a control. A vacuole is seen in the

BafA1 treated parental cell. Bottom, parental and ΔATG5 promastigotes expressing mNG::ATG8 as an autophagosome marker. Arrows indicate the ATG8 signal DMSO-treated cells and in the vacuole (enlarged autolysosome) of BafA1-treated cells. ΔATG5 cells show no ATG8 puncta and no enlarged autolysosomes after 3 days of BafA1 treatment. Magenta: DNA, scale bars are 5 µm. (**E**) Fluorescence micrographs of pHLuorin2::CPA expressing cells treated with BafA1 (400 nM) or DMSO (control) for 3 days. pHLuorin2::CPA positive enlarged vacuoles are seen in the BafA1 treated cells (arrows). Scale bars: 10 µm.

**Supplementary Figure 4: Diagnostic PCR for $V_1$E knockout verification**

Agarose gels showing diagnostic PCRs to test for the presence of the v-ATPase subunit $V_1$E gene, to verify the $V_1$E knockout in clonal cell lines tagged with (**A**) mNG::ATG8, (**B**) mNG::CPA, (**C**) pHLuorin2::CPA. First lane: Gene Ruler 100 bp DNA ladder (Invitrogen). The IFT88 gene was amplified as a control to test for the presence of genomic DNA in the samples. Amplification from the parental gDNA (C9T7) is a positive control for the amplification of $V_1$E.

**Supplementary Figure 5: Diagnostic PCR amplification of ATG5 for knockout verification**

Agarose gels showing diagnostic PCRs to test for the presence of the ATG5 gene, to verify the knockout in (**A**) the parental C9T7 cell line, (**B**) mNG::ATG8 tagged promastigotes. First lane: Gene Ruler 100 bp DNA ladder (Invitrogen). The IFT88 gene was amplified as a control to test for the presence of genomic DNA in the samples. Amplification from the parental gDNA (C9T7) is a positive control for the amplification of ATG5.

**Supplementary Table 1: Localisation summary**

v-ATPase subunit localizations and summary of $V_1$E co-localisation analyses.

**Supplementary Table 2: KO cell lines and primers**

Summary of knockout (KO) cell lines and ORF primers for KO validation.

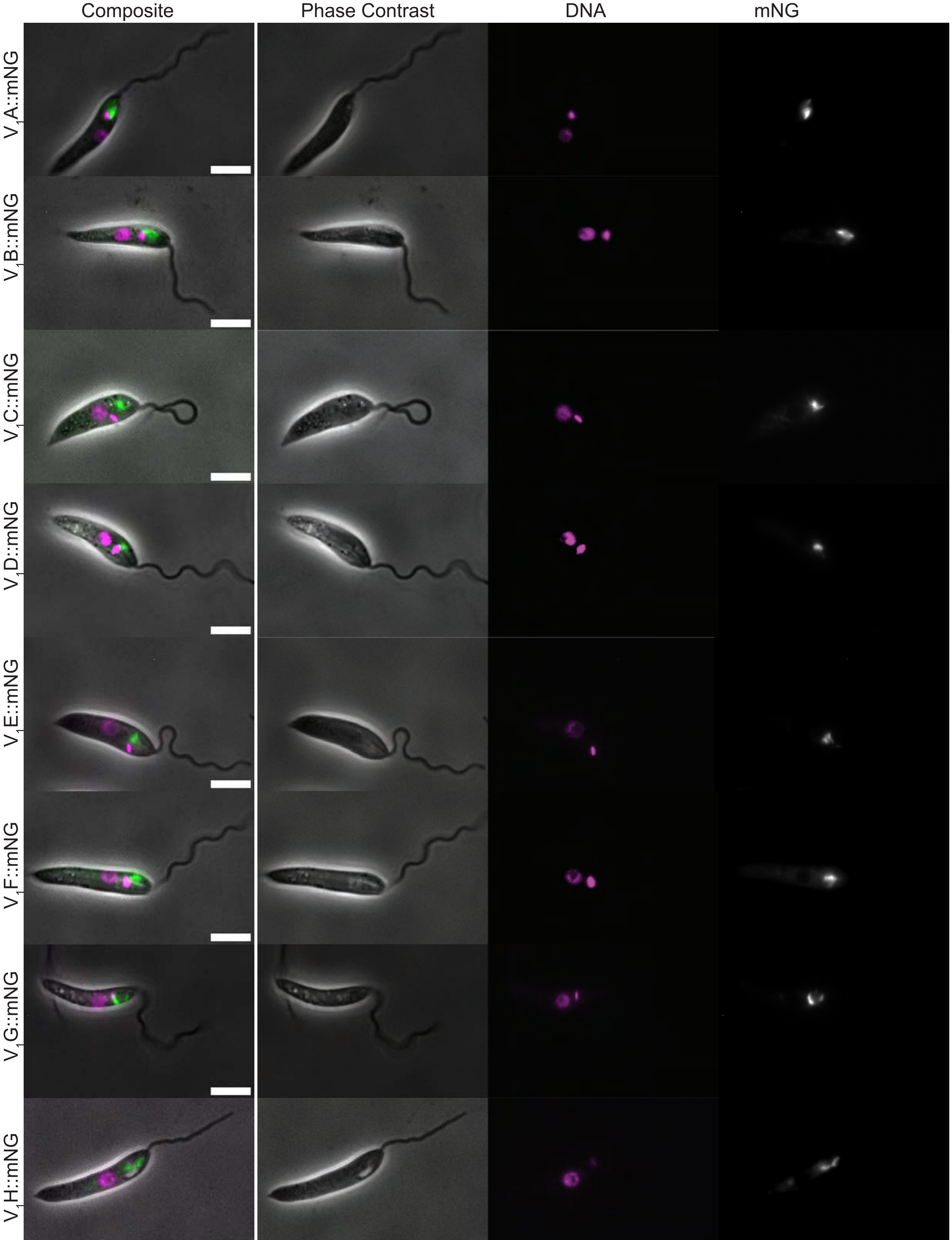


**Supplementary Figure 1A**

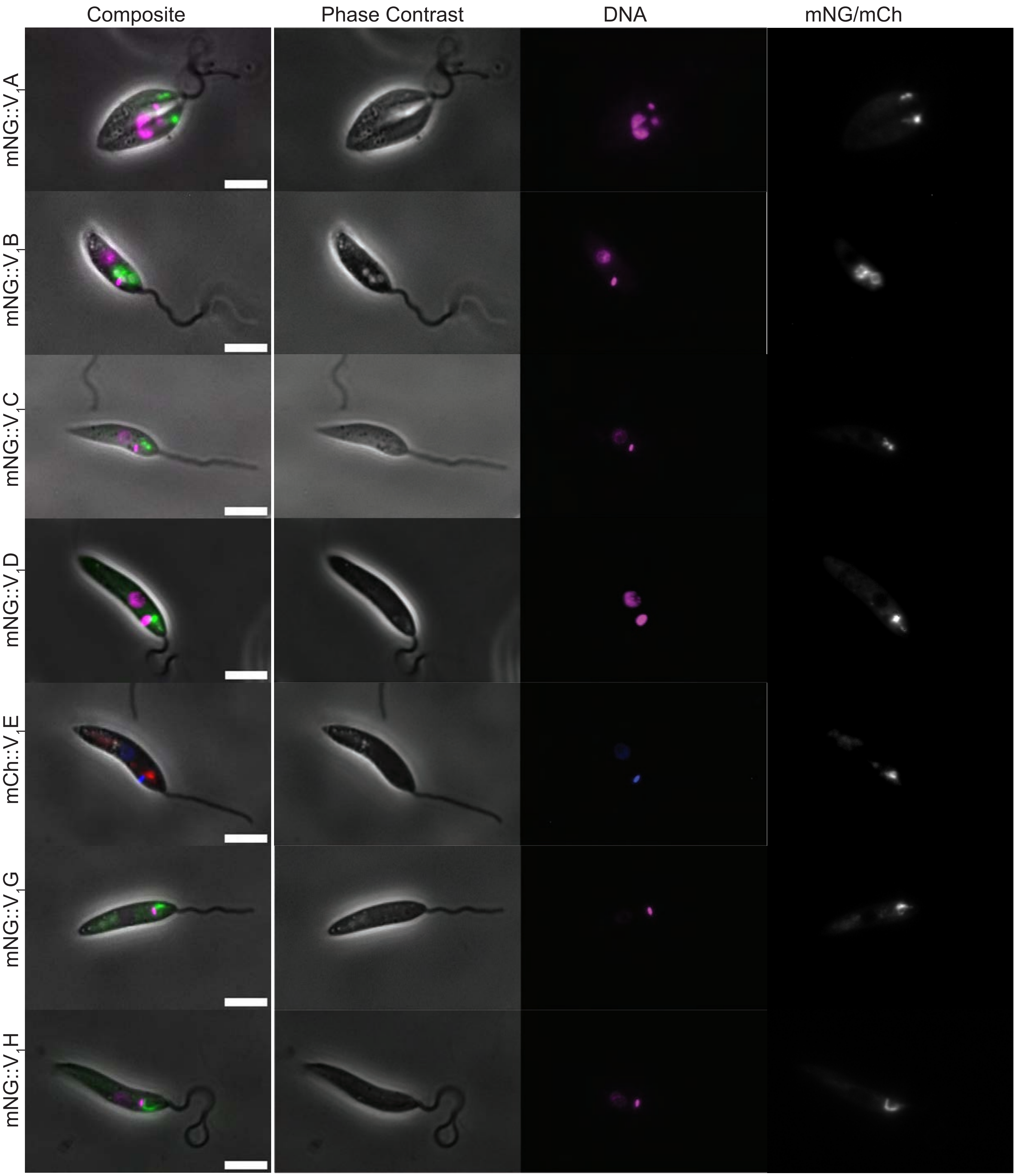


**Supplementary Figure 1B**

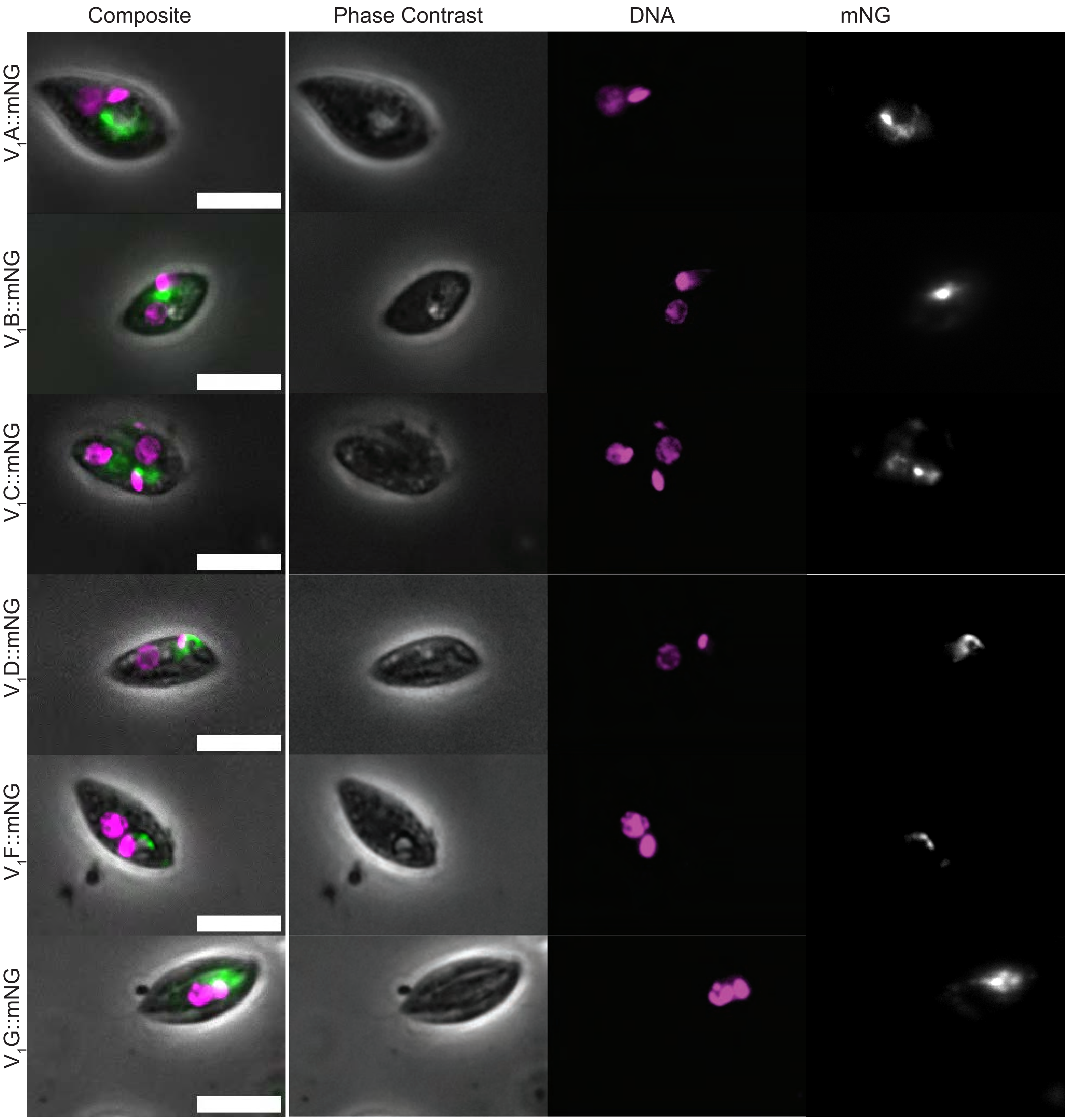


**Supplementary Figure 1C**

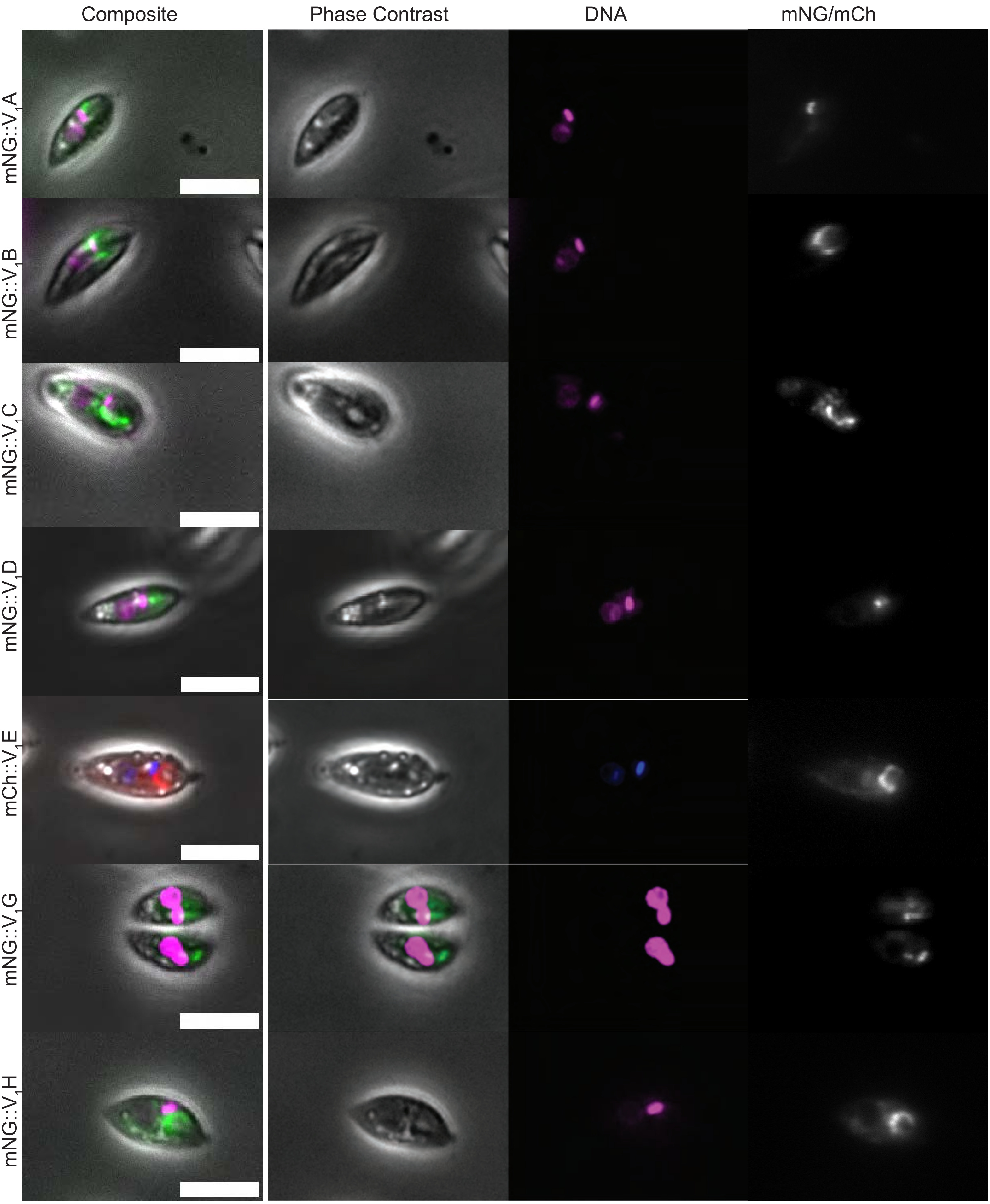


**Supplementary Figure 1D**

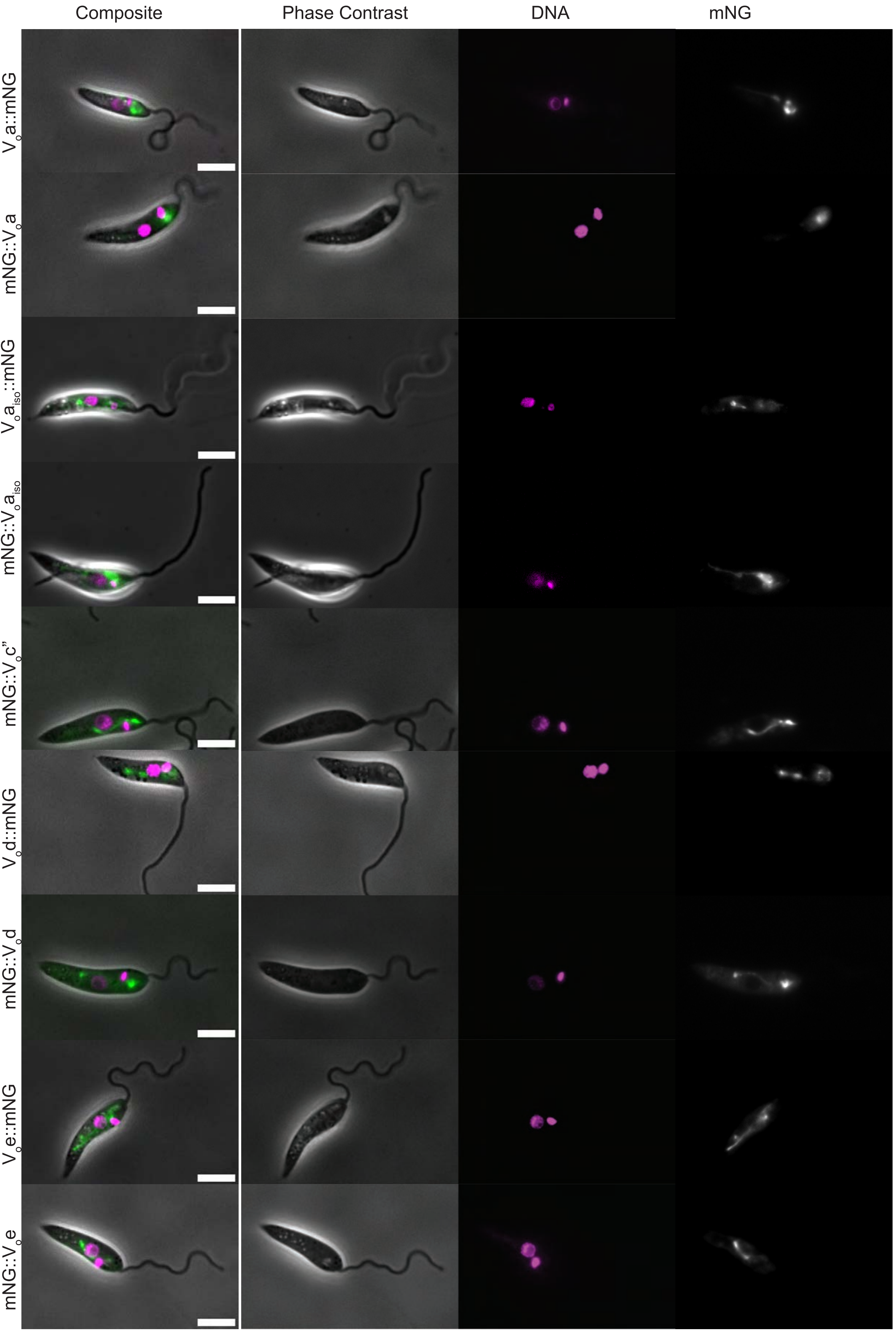


**Supplementary Figure 1E**

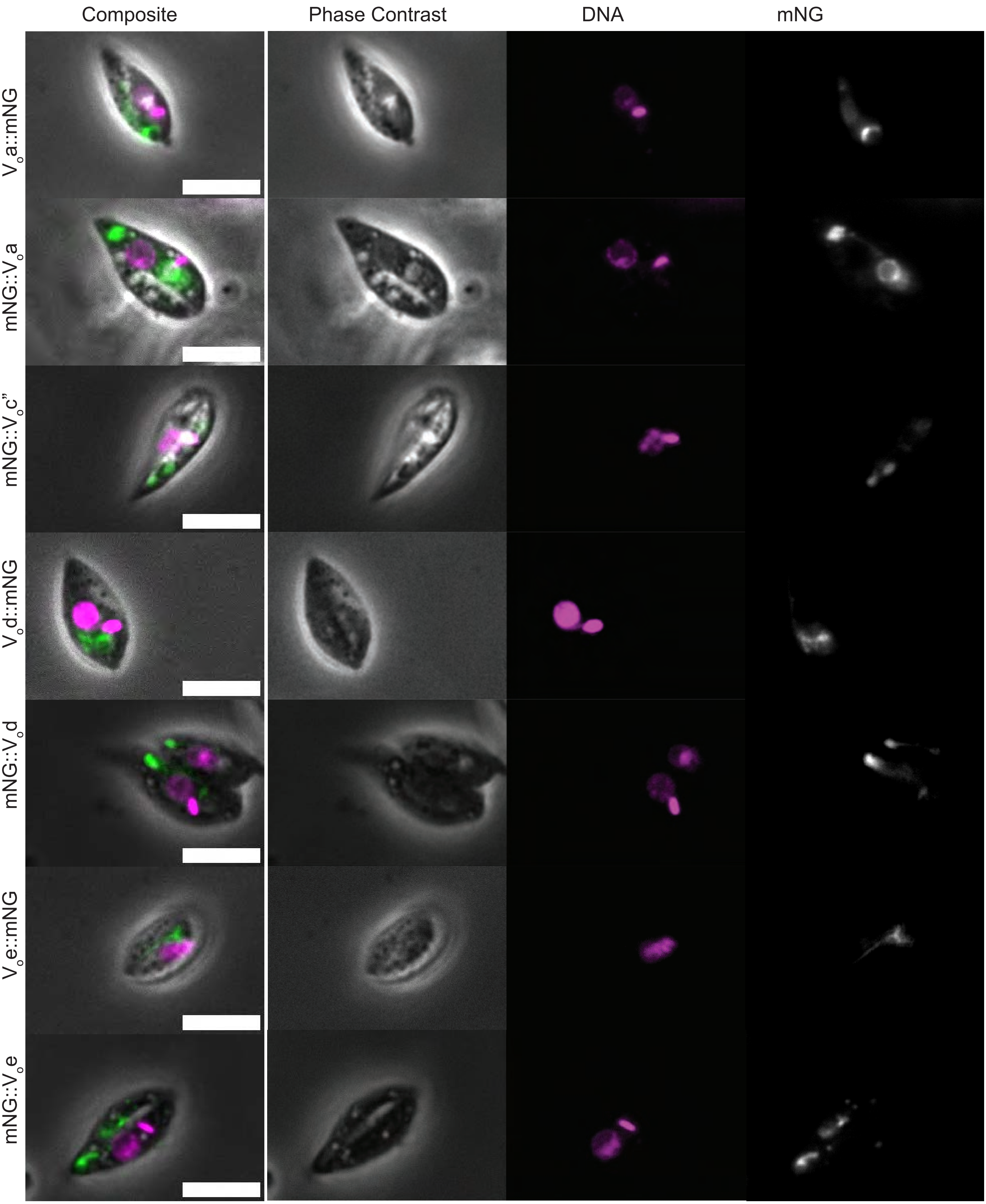


**Supplementary Figure 1F**

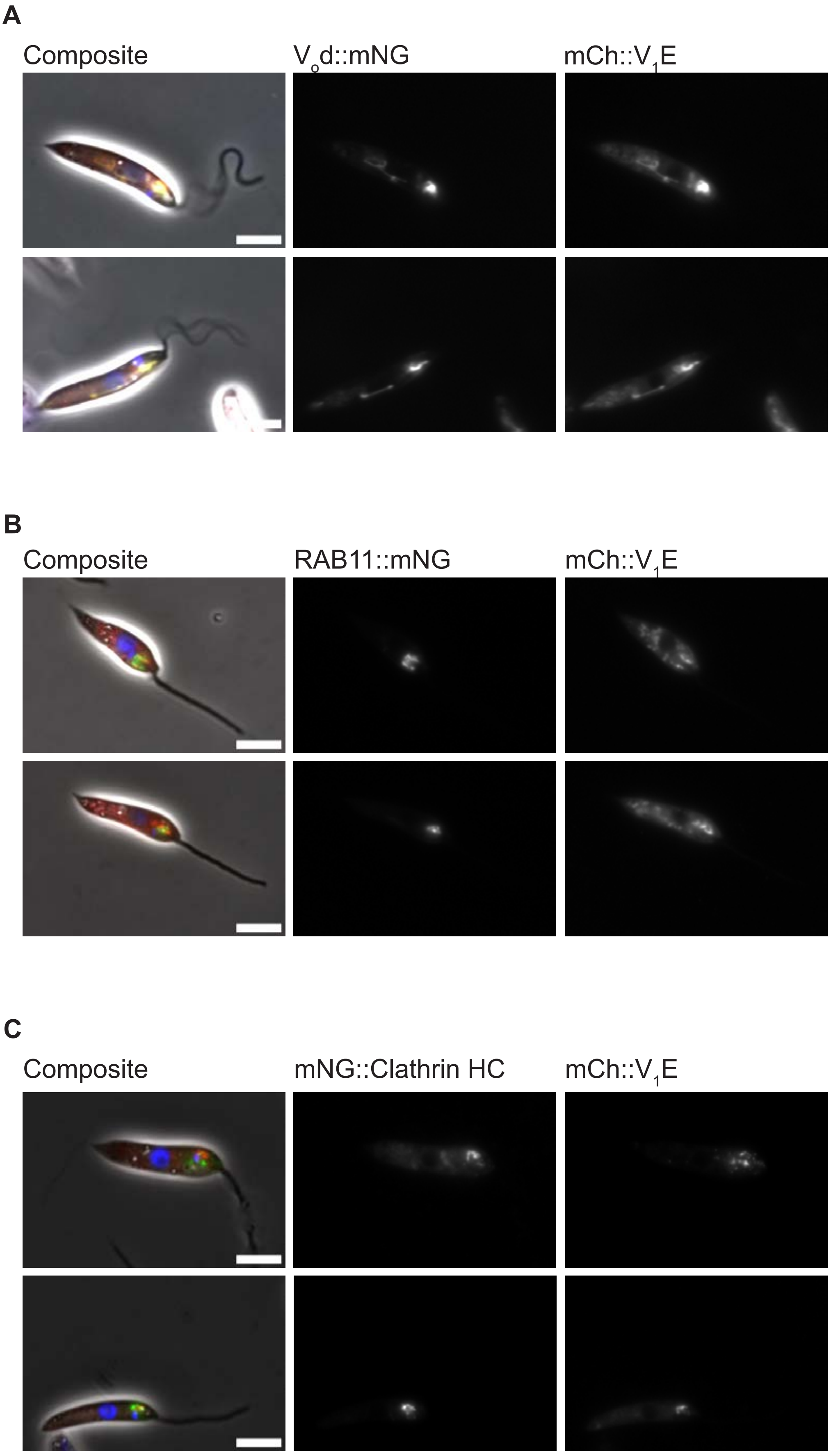


**Supplementary Figure 2**

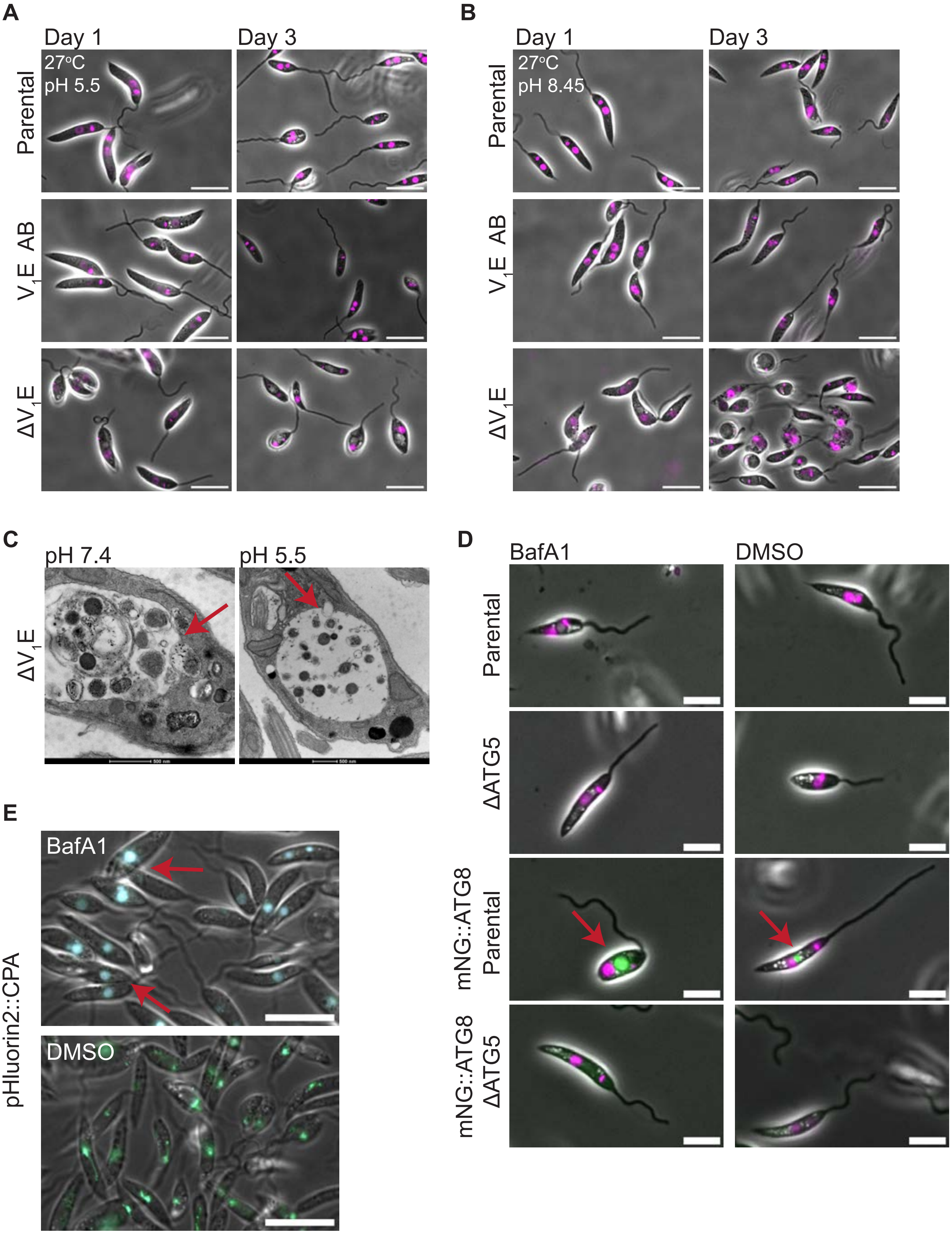


**Supplementary Figure 3**

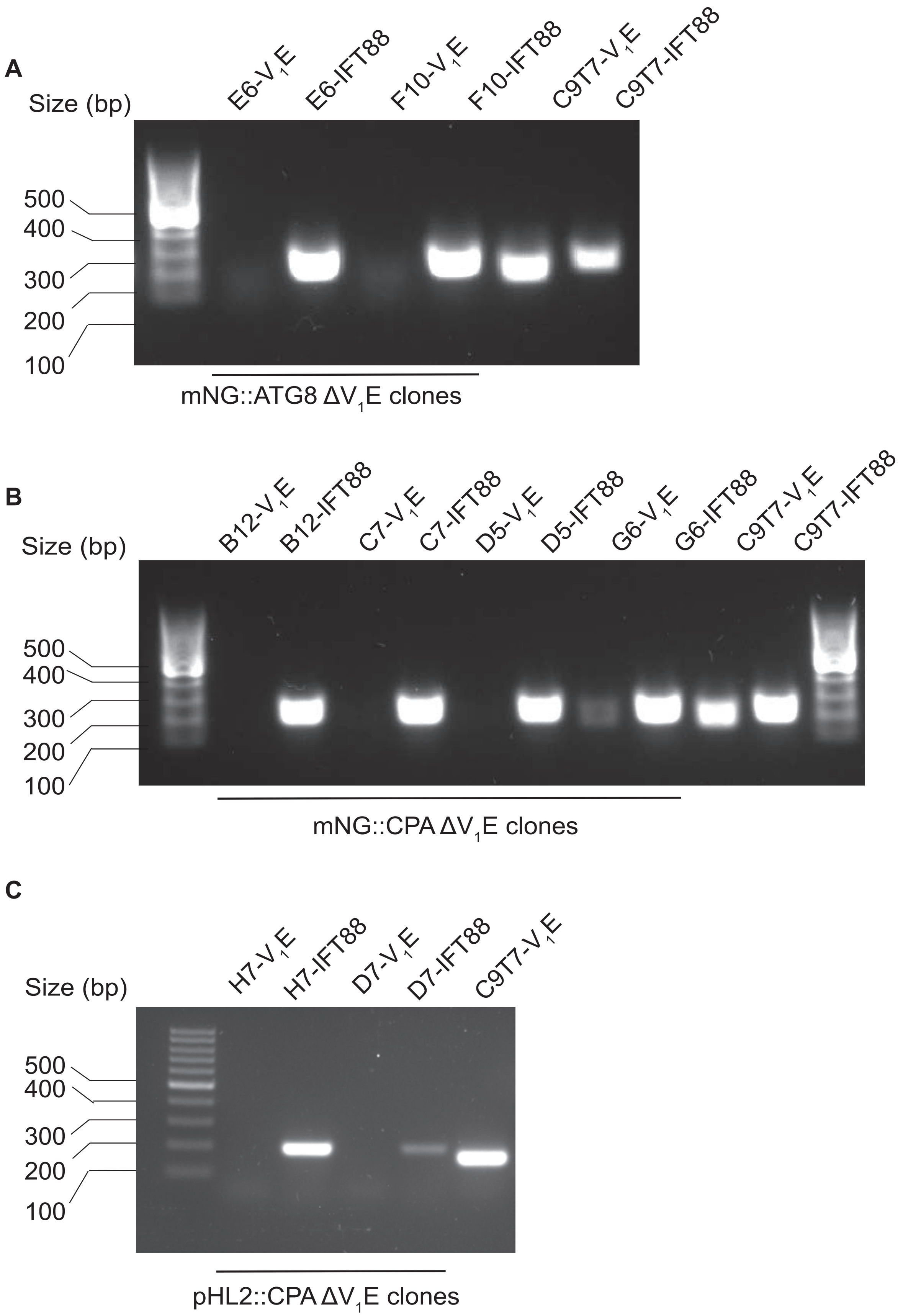


**Supplementary Figure 4**

**A**

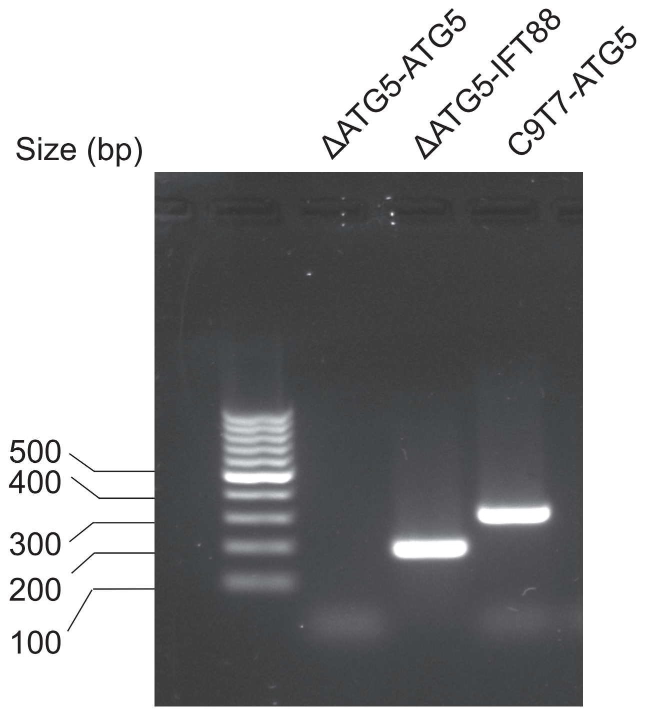


**B**

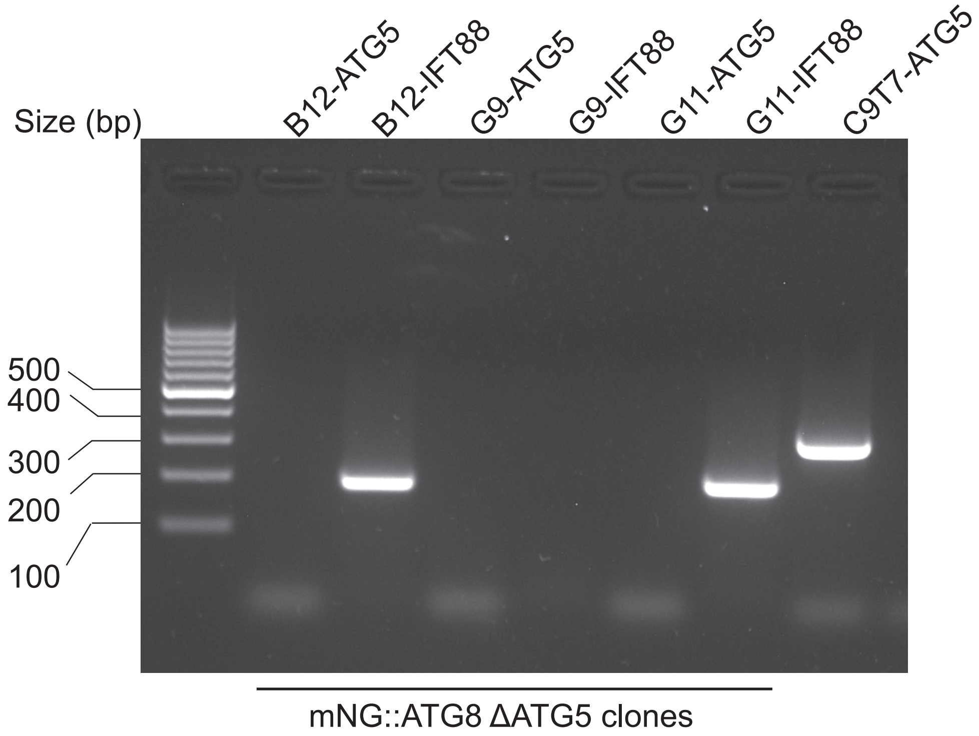


**Supplementary Figure 5**
